# Co-repressor AtSDR4L and paralog regulate hormonal and hypoxia responses in multiple Arabidopsis seed compartments

**DOI:** 10.1101/2024.11.29.626136

**Authors:** Bailan Lu, Dongeun Go, Jiayi Shan, Liang Song

## Abstract

Development is a series of decision-making events. Success of seed plants at individual and population levels strongly depends on the timing of germination and the rapidness of seedling establishment. *Arabidopsis thaliana* SEED DORMANCY 4-LIKE (AtSDR4L) and its paralog Dynamic Influencer of Gene Expression 2 (DIG2) are transcriptional co-repressors that promote seed-to-seedling phase transition. Their regulatory roles in promoting germination at the temporal and tissue-specific scales remain elusive. We show that strong germination arrest of *Atsdr4l dig2* is alleviated by ABA antagonists. Isolated mutant embryos develop faster than intact seeds, but still exhibit delayed growth. *Atsdr4l dig2* seeds show extensive changes in gene expression in both the seed coat and the embryo, with a subset of the genes differentially expressed tissue-specifically. MIKC-type MADS-box genes are the top-enriched transcription factor family among up-regulated genes in both seed compartments of *Atsdr4l dig2*, and *AGAMOUS-LIKE44* (*AGL44*) is a dirct target of both AtSDR4L and DIG2. Many hormonal genes and hypoxia-responsive genes are misregulated in the double mutant seeds, accompanied by an over-accumulation of abscisic acid, auxin, their derivatives, as well as the immediate precursor of ethylene. Together, these results provide new mechanistic insights into how AtSDR4L and DIG2 work in concert to coordinate multiple pathways and prepare seeds for germination.

## Introduction

Seed-to-seedling transition involves break of dormancy, germination, and seedling establishment. During seed maturation, dormancy is acquired to prevent germination in otherwise favorable conditions. Being an adaptive trait, dormancy reflects the interplay between genetic and environmental factors. The strength and types of the dormancy vary by species, ecotypes, and are influenced by environmental and nutrient cues such as temperature, light, and nitrate (Baskin and Baskin, 2004; Iwasaki et al., 2022; Sajeev et al., 2024). Physiological dormancy is the best studied among five main types of seed dormancy (Baskin and Baskin, 2020). Arabidopsis seeds have non-deep physiological dormancy, characterized by a fully developed embryo and the ability to break dormancy through dry storage (after-ripening). Arabidopsis seed coat, including both the testa and the endosperm, plays important roles in dormancy (Debeaujon et al., 2000; Finch-Savage and Leubner-Metzger, 2006). Removal of the seed coat or the endosperm de-arrest embryo growth (Bethke et al., 2007; Lee et al., 2010).

Dormancy in Arabidopsis is maintained by high abscisic acid (ABA) levels upon imbibition as well as ABA-independent mechanisms (Iwasaki et al., 2022), whereas germination is promoted by elevated ratio of gibberellins (GA) to ABA (Kucera et al., 2005). The level of ABA is the outcome of ABA biosynthesis and inactivation. Several *9-cis-epoxycarotenoid dioxygenase* (*NCED*) genes, encoding enzymes that catalyze a committed step of ABA biosynthesis, are expressed with comprehensive spatiotemporal specificity in various compartments of developing and maturing seeds (Tan et al., 2003; Lefebvre et al., 2006; Frey et al., 2012). The level of *CYP707A2*, encoding a key enzyme for ABA catabolism, increases rapidly upon imbibition (Kushiro et al., 2004). Single mutant of *cyp707a2* and its higher-order mutants with other ABA 8-hydroxylases such as *cyp707a1* and *cyp707a3* over-accumulate ABA and are more dormant (Okamoto et al., 2006). Furthermore, the distribution of ABA in seeds is also determined by transport from the endosperm to the embryo by ABC transporters (Kang et al., 2015). GA level is low in dry seeds and increases in response to cold stratification or imbibition (Weitbrecht et al., 2011). Increase in GA enables cell wall loosening in both the endosperm and the embryo, and promotes elongation of the lower hypocotyl. Consequently, the rupture of the endosperm and protrusion of the radicle complete germination. Many other hormones, such as auxin, ethylene, jasmonates, and brassinosteroids modulate the balance between ABA and GA through crosstalk at the hormone biosynthesis and signaling steps to regulate the transition from seed to seedlings (Gazzarrini and Tsai, 2015; Skubacz and Daszkowska-Golec, 2017). Regulators such as DELAY OF GERMINATION 1 (DOG1), MOTHER OF FT AND TFL1 (MFT), ABA INSENSITIVE5 (ABI5), and RGA-LIKE 2 (RGL2) integrate light, temperature, and hormonal cues to modulate dormancy and germination (Graeber et al., 2014; Yang et al., 2020b).

*Arabidopsis thaliana SEED DORMANCY 4-LIKE* (*AtSDR4L*) encodes a transcriptional co-repressor that promotes seed-to-seedling transition (Cao et al., 2020; Liu et al., 2020; Wu et al., 2022; Zheng et al., 2022). AtSDR4L physically interacts with Polycomb Repressive Complex (PRC)-interacting transcription factor (TF) VIVIPAROUS1/ABI3-LIKE2 (VAL2) (Lu et al., 2024). AtSDR4L and VAL2 bind to the upstream of *LEAFY COTYLEDON1* (*LEC1*), *ABA INSENSITIVE3* (*ABI3*), its own gene and a few *AtSDR4L* paralogs (Lu et al., 2024). The binding is often associated with reduced level of repressive histone mark H3K27me3 at these loci (Lu et al., 2024). LEC1, ABI3, along with FUSCA3 (FUS3), and LEC2, are collectively known as LAFL master TFs that regulate seed development and their expression need to be turned off during seed-to-seedling transition (Gazzarrini and Song, 2024). All *LAFL* genes are de-repressed in *Atsdr4l* loss-of-function seedlings (Wu et al., 2022; Zheng et al., 2022). Defects in seed-to-seedling transition are exacerbated when both *AtSDR4L* and its paralogs are mutated, exhibiting severe germination arrest and extensive accumulation of seed storage compounds in seedlings (Zheng et al., 2022). The molecular mechanism of strong seed germination arrest has been attributed to elevated expression of *NCED* (Liu et al., 2020; Zheng et al., 2022). However, tissue-specific regulation of germination in the *Atsdr4l* mutants hasn’t been fully elucidated.

In this study, we examined how AtSDR4L and its paralogs promote germination and subsequent seed-to-seedling transition in the seed coat and the embryo by testing the response to ABA antagonists, seed coat bedding assays, seed compartment-specific transcriptome profiling, and comprehensive hormone analysis. We revealed an extensive and sophisticated interplay of hormonal, transcriptional, and hypoxic responses and provided potential mechanistic explanations for the elevated germination arrest through the derepression of the *AGAMOUS-LIKE* (*AGL*) TFs and *ABI5*, over-accumulation of ABA and auxin, and increased hypoxia responses in *Atsdr4l dig2*.

## Materials and Methods

### Plant materials and growth conditions

All wild-type control and mutant plants used in this study are from Columbia-0 (Col-0) background. *Atsdr4l-5* deletion mutant was generated by CRISPR-Cas9 in the previous study (Lu et al., 2024), and insertional mutants for *DIG1* and *DIG2* were obtained from SALK T-DNA collection (Alonso et al., 2003). The single mutants of *dig1* (SALK_128578) and *dig2* (SALK_057406) were first crossed to create their double mutant, and *Atsdr4l-5* was subsequently introduced by crossing with *dig1* and *dig2*. *Atsdr4l-5 dig2* and *Atsdr4l-5 dig1* were segregated from F3 progeny of the cross and confirmed using PCR genotyping primers listed in Table S1. Col-0 wildtype and the higher-order mutant seeds were surface sterilized and plated on 1X Linsmaier & Skoog (LS) medium (LSP03, Caisson Labs, Smithfield, UT, USA) containing 0.7% (w/v) agar (A111, PhytoTech Labs, Lenexa, KS, USA). Seeds were cold stratified for 3 days at 4 °C and grown on plates for 2 to 4 weeks, then transferred to soil and grown under a “long-day” condition (16 h light at 23 °C / 8 h dark at 18 °C). *Agrobacterium tumefaciens* strain GV3101 was used to produce the transgenic plants (in *Atsdr4l-5 dig2* background) expressing *AtSDR4Lpro::3XHA-AtSDR4L-3XFLAG* or *DIG2pro::3XHA-DIG2-3XFLAG* through the floral dip method (Clough and Bent, 1998). Selection of the transgenic lines was carried out on 1X LS medium containing 25 µg/mL Hygromycin B (H-270-1, GoldBio, USA). T2 seeds for the examination of complementation were collected when siliques turned yellow-to-brown. After 12 days of storage at room temperature, embryos were isolated after briefly imbibing seeds for 30 minutes, and directly bedded on 1X LS medium with a mesh.

### Plasmid construction

The vectors for complementing the *Atsdr4l-5 dig2* mutants were generated as described in the previous study (Lu et al., 2024). Primers listed in TableS1 were used to construct the vector of *AtSDR4Lpro::3xHA-AtSDR4L-3xFLAG* by amplifying the promoter and the fragment containing *3xHA-AtSDR4L-3xFLAG* from an existing plasmid published in the previous study (Lu et al., 2024). *DIG2pro::3xHA-DIG2-3xFLAG* was generated following the similar procedure using primers listed in Table S1. In brief, 2781-bp promoter and the coding sequence (without stop codon) of *DIG2*, N-terminal 3xHA and C-terminal 3xFLAG tags were cloned into the linearized pCAMBIA1300 vector after EcoRI and PstI digests using the NEBuilder® HiFi DNA Assembly Master Mix following the manufacturer’s instructions (E2621S, NEB, Ipswich, MA, USA).

Cloning of the vectors pCAMBIA1300-AtSDR4Lpro::3xHA-AtSDR4L-3xFLAG and pCAMBIA1300-DIG2pro::3xHA-DIG2-3xFLAG was performed using *Escherichia coli* competent cells of NEB 10-beta strain (C3019H, NEB, Ipswich, MA, USA) and One Shot™ Mach1™ T1 strain (C862003, Thermo Fisher Scientific, Waltham, MA, USA). The plasmids confirmed by colony PCR and Sanger sequencing were transformed into the *A. tumefaciens* competent cells GV3101 by electroporation, and transfected GV3101 cells were further verified by colony PCR. Transgene insertion in the T1 complementation plants were confirmed by PCR genotyping.

### Germination assays

The physiological characterization of the single and higher-order mutants was carried out using 3 biological replicates for each genotype. Seeds were collected at maturity at roughly 18 days after pollination (DAP), when siliques turned yellow-to-brown, and after-ripened for 2 weeks. After surface sterilization, seeds were directly sowed, or stratified at 4°C for 3 days and then sowed on 1X LS (LSP03, Caisson Labs, Smithfield, UT, USA) medium with 0.7% (w/v) agar, and supplemented with the following either hormone or hormone antagonists or both: 20 µM gibberellins (GA4 + GA7) (G026-1GM, Caisson Labs, Smithfield, UT, USA), 50 µM antabactin (Vaidya et al., 2021), 50 µM ABA-1019 (Diddi et al., 2021), 20 µM GA4 + GA7 with 50 µM antabactin, and 20 µM GA4 + GA7 with 50 µM ABA-1019. DMSO was used as the solvent and included as the mock treatment. Plates were incubated under a “long-day” condition (16 h light at 23°C / 8 h dark at 17°C), and the percentage of germination was scored daily for up to 7 days after imbibition (DAI). The percentage of germination was compared across all groups using ANOVA test followed by Tukey’s HSD to determine the statistical significance between each pair of treatments. For ABA and D-mannitol treatments, embryos were isolated from 3-week after-ripened seeds that were cold stratified for 3 days at 4°C. In each biological replicate, six embryos were directly sowed on 1X LS medium with 0.7% (w/v) agar containing either ABA (A-050-1, Gold Biotechnology, St Louis, MO, USA) at 0.1 µM or D-mannitol (A14030, Alfa Aesar, Ward Hill, MA, USA) at 300 mM. The responsiveness of the *Atsdr4l-5 dig2* mutant and Col-0 to ACC was examined by cold stratification of 3-week after-ripened seeds for 3 days, and then plating on 1X LS medium with 0.7% (w/v) agar supplemented with 50 µM ACC (J65118.MD, Thermo Fisher Scientific, Waltham, MA, USA). Germination rate was scored for up to 4 days. One-tailed Student’s t-test was used to compute the statistical significance in the difference between mock and ACC-treated Col-0 and *Atsdr4l-5 dig2* mutant seeds.

### Seed coat bedding assay

Seed coat bedding assay was performed following the protocol by Lee and Lopez-Molina (Lee and Lopez-Molina, 2013). In brief, seeds were harvested at maturity at roughly 18 DAP from yellow-to-brown siliques and after-ripened for 2 days prior to surface sterilization and cold stratification for 3 days at 4°C. In each biological replicate, ten to twelve embryos were isolated from the seed coats and sowed directly either on the 80-micron nylon mesh or on 30 to 40 seed coat beddings on the surface of 1X LSmedium with 0.7% (w/v) agar. Plates were incubated under 8 h dark, 17°C / 16 h light, 23°C for 4 days to observe the effect of different bedding strata on seedling establishment.

### Image acquisition and processing

Images of the embryos grown on mannitol and ABA media were captured using Leica M205 FCA microscope with objective lens/numerical aperture (NA) set at 1x/0.04 and magnification set at 0.82. Samples from the seed-coat bedding assay were seen with objective lens/numerical aperture (NA) at 1x/0.03 and magnification at 0.64. Acquired images were processed with Fiji software v1.54k.

### Preparation of seed compartment RNA-seq library

Seeds collected at maturity from ∼ 18 DAP, yellow-to-brown siliques were after-ripened for 2 days. Batch dissection of seeds and collection of embryos and seed coat were performed following the procedure described by Iwasaki and Lopez-Molina (Iwasaki and Lopez-Molina, 2021). In brief, seeds were imbibed in water for thirty minutes at room temperature before dissection. For each biological replicate, approximately 50 to 100 seeds were gently pressed against two microscope glass slides to break the seed coats. Seed coat (containing endosperm) and intact embryos were manually separated using syringe needles and fine forceps and preserved in RNAlater buffer (AM7020, Thermo Fisher Scientific, Waltham, MA, USA) in a microcentrifuge tube until all samples have been collected within two hours of imbibition. RNAlater buffer was vacated using a syringe before samples were snap frozen in liquid nitrogen. Extraction of total RNA was carried out following the procedure as described in the previous studies (Wu et al., 2022; Lu et al., 2024). Briefly, total RNA was isolated using Spectrum™ Plant Total RNA Kit (STRN50, MilliporeSigma, Burlington, MA, USA), followed by the removal of genomic DNA by DNase I treatment (AM1907, TURBO DNA-free™ Kit, Thermo Fisher Scientific, Waltham, MA, USA). Total RNA from embryo and endosperm-containing seed coat of Col-0 and *Atsdr4l-5 dig2* seeds (same stage and storage conditions as samples for seed coat bedding and seed dissection) were used to make cDNA libraries with three biological replicates each. NEBNext® Ultra II Directional RNA Library Prep Kit for Illumina® (E7760S, NEB, Ipswich, MA, USA) with NEBNext Poly(A) mRNA Magnetic Isolation Module (NEB #E7490) and Unique Dual Index UMI Adaptors RNA Set 1 (E7416S, NEB, Ipswich, MA, USA) were used for library preparation following the manufacturer’s instructions. Sequencing was carried out at Novogene with NovaSeq X Plus PE150.

### RNA-seq analysis

Mapping and transcript quantification: raw sequencing reads from the tissue-specific RNA-seq data were preprocessed with fastp v0.12.4 for adaptor trimming (Chen et al., 2018), and subsequently aligned to the Arabidopsis TAIR10 reference genome with Araport11 annotation using STAR 2.7.9a and the arguments “*--runMode alignReads --outFilterMultimapNmax 10 -- outFilterMismatchNoverLmax 0.05 --quantMode GeneCounts --outSAMtype BAM SortedByCoordinate --outReadsUnmapped Fastx*” (Dobin et al., 2013). FPKM quantification of transcripts from RNA-seq was determined using salmon v1.10.1 tool (Patro et al., 2017). Preprocessed RNA-seq reads from fastp v0.12.4 were quantified with salmon under “*--l A*” parameter for the auto library type detection. Salmon transcript quantification data was loaded using tximport v1.30.0 (Soneson et al., 2015). *Arabidopsis thaliana* TAIR10.50 GTF annotation file was then processed with GenomicFeatures v1.54.4 to make an annotated transcript database, to which extracted transcript IDs and corresponding gene IDs were mapped for transcript-to-gene mapping of FPKM values (Lawrence et al., 2013).

Differential gene expression analysis: read count data were extracted from the “ReadsPerGene” output file and DEGs were computed using Limma v3.50.3 (Ritchie et al., 2015) in RStudio v2021.09.2+382 (The R Development Core Team, 2020). Genes were filtered by counts per million (CPM) values greater than 1 in at least 25% of the samples. Library size was normalized using “*Trimmed Means of M-values (TMM)*” method in edgeR v3.36.0 function “*calcNormFactors*” (Robinson et al., 2010). Principal component analysis was performed using “plotMDS” function in Limma v3.50.3 using all detected genes after read count filtering, with the parameter “gene.selection” set to “common” (Ritchie et al., 2015). Pair-wise correlation among embryo and seed coat samples were computed using log_2_(CPM+1) values and graphed by pheatmap v1.0.12 (Kolde, 2019) with viridis colour palettes (Garnier, 2018). Log_2_ fold changes (LFC) and FDR-adjusted p-values were computed by contrasting the mutants with the wild-type samples. Genes with LFC > 0.5 and < -0.5 and adj.P.Val < 0.05 were considered significantly up- and down-regulated. DEGs shared among all the mutants were extracted and intersected using tidyverse v1.3.2 (Wickham et al., 2019).

Enrichment analysis: the UpSet plots were generated using the “*UpSet*” function in ComplexHeatmap v2.10.0, with “*intersection*” mode specified in the combination matrix (Gu et al., 2016). A list of 49 genes (core 49) that are induced by hypoxia was obtained from existing literature (Mustroph et al., 2009) and intersected with the significantly up- and down-regulated genes in *Atsdr4l-5 dig2* seed compartments (LFC > 0.5 or LFC < -0.5 and adj.P.Val < 0.05). The p-values for the overlap between each DEG list and core 49 were computed by GeneOverlap v1.30.0 (Shen, 2017). The total number of all expressed genes in the tissue-specific RNA-seq (after low read count filtering) was used as the genomic background. Up-regulated genes in *Atsdr4l-5 dig2* (same filtering criteria as UpSet plot) were used for transcription factor (TF) family enrichment analysis. TF family annotation was obtained from Plant Transcription Factor Database and AGRIS Arabidopsis transcription factor database (Davuluri et al., 2003; Tian et al., 2019), and the two lists were combined and filtered for unique genes before intersecting with the expressed genes in RNA-seq. All TFs detected in the tissue-specific RNA-seq experiment after filtering out genes with low read count were used as the background list for the hypergeometric test with the “*enricher*” function in clusterProfiler v4.2.2 (Yu et al., 2012). The statistics for the top enriched family in both seed compartments were presented. Gene symbols for the up-regulated MIKC MADS members were converted from their TAIR IDs using org.At.tair.db v3.14.0 (Carlson, 2017). Gene set enrichment analysis (GSEA) of the gene ontology (biological process) terms was performed using the “*gseGO*” function in clusterProfiler v4.2.2 (Yu et al., 2012) with 100 permutations and org.At.tair.db 3.14.0 annotations (Carlson, 2017). Significantly up-regulated and down-regulated genes in the mutant embryo and seed coat were filtered by LFC > 0.5 and LFC < -0.5, respectively, and ranked by the absolute LFC values by descending order in the corresponding tissues as the input lists for GSEA. Differential expression data of ABA-responsive genes were obtained from GSE80565 (Song et al., 2016) for plotting with the DE results from tissue-specific RNA-seq. Genes near ChIP-seq peaks were annotated by ChIPpeakAnno v3.28.1 using Araport11 genome annotation (Zhu et al., 2010). The maximum distance of 3 kb was used to define the overlaps between peaks and genes. Genes with peaks of -log_10_[qValue] ≥ 10 were considered candidate targets of AtSDR4L or DIG2. Unique TAIR IDs were kept and intersected with the RNA-seq data to generate the UpSet plots using ComplexHeatmap v2.10.0 (Gu et al., 2016). GeneOverlap v1.30.0 was used to compute the p-values of overlaps between each DEG list and the ChIP-seq targets (Shen, 2017).

Visualization: tiled data files (TDF) were generated from BAM files using igvtools v2.5.3 with the parameters “*--includeDuplicates -w 10 -z 5*” to specify the maximum zoom level at 5 for precomputing and window size at 10 bp for averaging coverage (Robinson et al., 2011). ChIP-seq data for 3xHA-AtSDR4L-3xFLAG (1-day old seedling) and GFP-DIG2 (4-day old seedling) were obtained from GSE246997 and PRJNA319317 (Song et al., 2016; Lu et al., 2024), respectively, and TDF files were geneared using the same parameters as described previously. TDF files were loaded on Integrative Genomics Viewer v2.14.0 (Robinson et al., 2011) with Araport11 annotations (Cheng et al., 2017) to visualize the transcript abundance and binding evidence at individual loci.

### RT-qPCR

To validate the expression of hypoxia-related genes, total RNA was isolated from mature seeds harvested at ∼ 20 DAP and after-ripened for 2 days. RNA extraction followed the same procedure as detailed in the “RNA-seq” section. For each sample, a total of 1000 ng total RNA was treated with DNase I (AM1907, TURBO DNA-free™ Kit, Thermo Fisher Scientific, Waltham, MA, USA) to remove genomic DNA. cDNA synthesis was carried out using Maxima H Minus Reverse Transcriptase following the manufacturer’s instructions (Thermo Fisher Scientific, Waltham, MA, USA, EP0752) and diluted by 20 folds before quantitative PCR conducted with the PowerUp™ SYBR™ Green Master Mix (Thermo Fisher Scientific, Waltham, MA, USA, A25742) in a QuantStudio™ 3 System (Thermo Fisher Scientific, Waltham, MA, USA) with 3 biological replicates. Fold change was computed by the ΔΔCT method with *ACTIN8* (AT1G49240) as the internal control.

### Phytohormone measurement

Hormone quantification was performed at the National Research Council Canada, Saskatoon (NRCC SK). Three sets of biological replicates, each containing approximately 55 mg of *Atsdr4l dig2* and 55 mg of Col-0 seeds harvested within two weeks since the siliques turn yellow, were used for hormone profiling. Analysis was performed on a UPLC/ESI-MS/MS utilizing a Waters ACQUITY UPLC system, equipped with a binary solvent delivery manager and a sample manager coupled to a Waters Micromass Quattro Premier XE quadrupole tandem mass spectrometer via a Z-spray interface. MassLynx™ and QuanLynx™ (Micromass, Manchester, UK) were used for data acquisition and data analysis. The procedure for quantitation of ABA, ABA catabolites, auxins, and gibberellins in samples of Arabidopsis seed tissue was performed using a modified procedure described in Lulsdorf et al. (Lulsdorf et al., 2013), while quantitation of ACC was performed using a modified procedure described in Chauvaux et al. (Chauvaux et al., 1997). A number of analyte standards namely DPA, ABA-GE, PA, 7’-OH-ABA, neoPA, trans-ABA and IAA-Glu were synthesized and prepared at the National Research Council of Canada, Saskatoon, SK, Canada; ABA, IAA-Leu, IAA-Ala, IAA-Asp, IAA, ACC, phenyl isothiocyanate (PITC), trifluoroacetic acid (TFA) and triethylamine were purchased from Sigma–Aldrich; GAs 1, 3, 4, 7, 8, 9, 19, 20, 24, 29, 44, and 53 were purchased from OlChemim Ltd. (Olomouc, Czech Republic). Deuterated forms of the hormones which were used as internal standards include: d3-DPA, d5-ABA-GE, d3-PA, d4-7’-OH-ABA, d3-neoPA, d4-ABA, d4-trans-ABA, d3-IAA-Leu, d3-IAA-Ala, d3-IAA-Asp, d3-IAA-Glu, and 13C4-IBA were synthesized and prepared at NRCC SK according to Abrams et al. (Abrams et al., 2003) and Zaharia et al. (Zaharia et al., 2005). The d5-IAA was purchased from Cambridge Isotope Laboratories (Andover, MA); 1-amino-[2,2,3,3-d4]-cylopropane-1-carboxylic acid (d4-ACC) was purchased from CDN isotopes (Point-Claire, QC, Canada); d3-dhZ, d3-dhZR, d5-Z-O-Glu, d6-iPR, d6-iP, 15N4-kinetin and d2-GAs 1, 3, 4, 7, 8, 9, 19, 20, 24, 29, 34, 44, 51 and 53 were purchased from OlChemim Ltd. (Olomouc, Czech Republic). The d4-ACC derivative PTH-d4-ACC was used as an internal standard. The deuterated forms of selected hormones used as recovery (external) standards were prepared and synthesized at NRCC SK. Calibration curves were created for all compounds of interest, including the ACC derivatives, phenylthiohydantoin-ACC (PTH-ACC) and PTH-d4-ACC. Quality control samples (QCs) were run along with the tissue samples. Hormone content was normalized against fresh tissue weight, and converted from ng / mg of samples to pmol / g according to the following molecular weights: ABA 264.32 g / mol, DPA 282.33 g / mol, ABAGE 426.46 g / mol, 7’OH-ABA 280.32 g / mol, IAA 175.18 g / mol, IAA-Asp 290.27 g / mol, IAA-Glu 304.30 g / mol, ACC 101.10 g / mol, t-ZR and c-ZR 351.36 g / mol. Statistical comparisons on the hormones and metabolites were performed using two-tailed Student’s t-test ith 3 replicates for each genotype.

## Results

### *Atsdr4l dig2* phenotypes are attenuated by ABA antagonists

Previously, multiple groups showed that *Atsdr4l* single and higher-order mutants exhibited delayed seed germination and defective seedling establishment (Cao et al., 2020; Liu et al., 2020; Wu et al., 2022; Zheng et al., 2022). We crossed *Atsdr4l-5*, a CRISPR/Cas9 segmental deletion line of *AtSDR4L* (Lu et al., 2024) with T-DNA insertion lines of *DIG* genes (Fig. S1A). Consistent with a previous study (Zheng et al., 2022), *Atsdr4l-5 dig2* seeds exhibit more severe germination arrest than the single mutant and *Atsdr4l-5 dig1* (Fig. 1, S1B). Therefore, we focus on *Atsdr4l-5 dig2* in this study to examine the temporal and spatial regulation of seed-to-seedling transition by AtSDR4L and its paralogs.

**Fig. 1.**
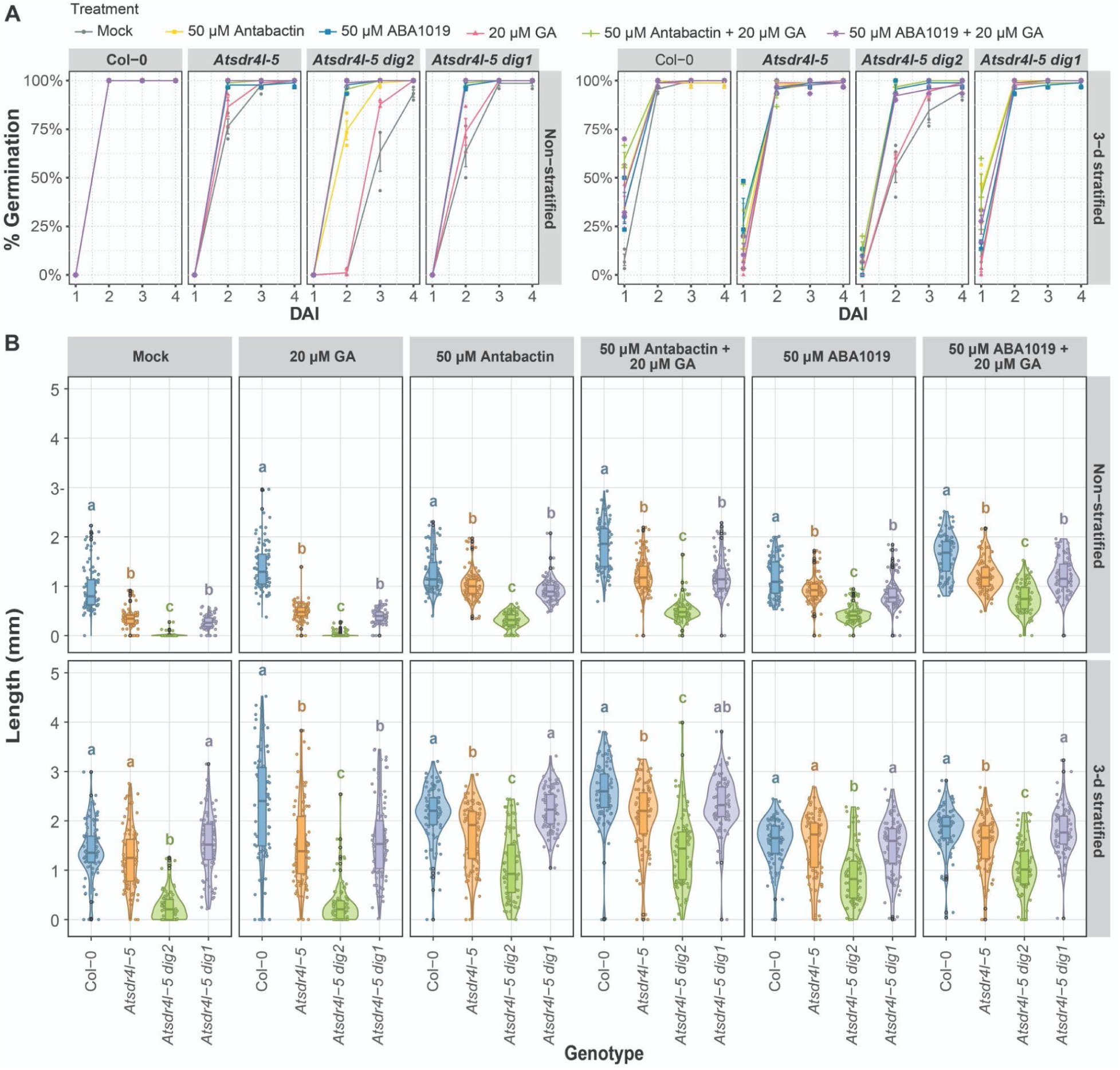
ABA antagonists promote germination and axis elongation of *Atsdr4l-5 dig2*. (A) Percentage of seed germination over days after imbibition (DAI) of non-stratified and 3-day cold stratified Col-0, *Atsdr4l-5*, *Atsdr4l-5 dig2* and *Atsdr4l-5 dig1* seeds on 1X LS 0.7 % (w/v) phytoagar medium with treatments of mock, 20 μM GA, 50 μM Antabactin, 50 μM Antabactin + 20 μM GA, 50 μM ABA1019, 50 μM ABA1019 + 20 μM GA. Error bars represent mean ± SEM (24 ≤ n ≤ 30, 3 biological replicates). (B) Quantification of axis length at 2 DAI in non-stratified and 3-day cold stratified Col-0, *Atsdr4l-5*, *Atsdr4l-5 dig2* and *Atsdr4l-5 dig1* seeds and seedlings grown on 1X LS 0.7 % (w/v) phytoagar medium. The black dots represent outliers. Different letters (*p* < 0.05) represent significant differences (24 ≤ n ≤ 30, 3 biological replicates, Kruskal-Wallis test, Dunn’s test).

Because ABA is a strong inhibitor of germination, we germinated wild-type and mutant seeds on plates supplemented with ABA antagonist antabactin (Vaidya et al., 2021) or antagonist / agonist ABA1019, formerly known as compound 7 (Diddi et al., 2021). At 50 µM, both compounds promote germination of *Atsdr4l-5* single and double mutants (Fig. 1, S1B-C). Since freshly harvested *Atsdr4l* seeds were reported to be unresponsive to GA (Cao *et al*., 2020), we applied GA in the germination assays with or without ABA analogues. Our results confirmed that GA alone on briefly after-ripened *Atsdr4l-5 dig2* seeds barely promotes germination (Fig. 1, S1B-C). In contrast, including GA in ABA antagonist treatment further improves seed germination and promotes embryo axis elongation (Fig. 1). These results suggest that ABA plays a primary role in the germination arrest of *Atsdr4l-5 dig2*.

### Germination arrest in *Atsdr4l dig2* is contributed by both the embryo and the seed coat

Because hormones regulate germination through both the endosperm and the embryo, we carried out a seed coat bedding assay (Lee et al., 2010) to determine which tissue causes the strong germination arrest of *Atsdr4l-5 dig2*. In the assay, dissected wild-type Col-0 and mutant embryos were placed on either Col-0 or mutant seed coat that contains the testa and the endosperm. We observed a clear difference between Col-0 and *Atsdr4l-5* single or double mutant starting from 2 days after imbibition (DAI) (Fig. S2). By 3 DAI, Col-0 embryos on all bedding strata and intact seeds were almost fully established as marked by expanded green cotyledons, elongated hypocotyls and roots with collet hairs, whereas *Atsdr4l-5* single mutant was established at a slower rate than Col-0 (Fig. 2A). By contrast, the majority of isolated *Atsdr4l-5 dig2* embryos remained arrested for axis elongation and collet hair development; cotyledon expansion and greening were very subtle and less synchronized (Fig. 2B). These defects were attenuated when the double mutant was complemented with native promoter-driven *DIG2* or *AtSDR4L* (Fig. S2C). Collectively, these results reveal that the embryos of *Atsdr4l-5* and *Atsdr4l-5 dig2* contribute substantially to the enhanced germination arrest. Seedling establishment in intact seeds is slower than corresponding dissected embryos in all genotypes, confirming the positive regulation of germination arrest by the seed coat through physical and physiological constraints (Fig. 2, S2).

**Fig. 2.**
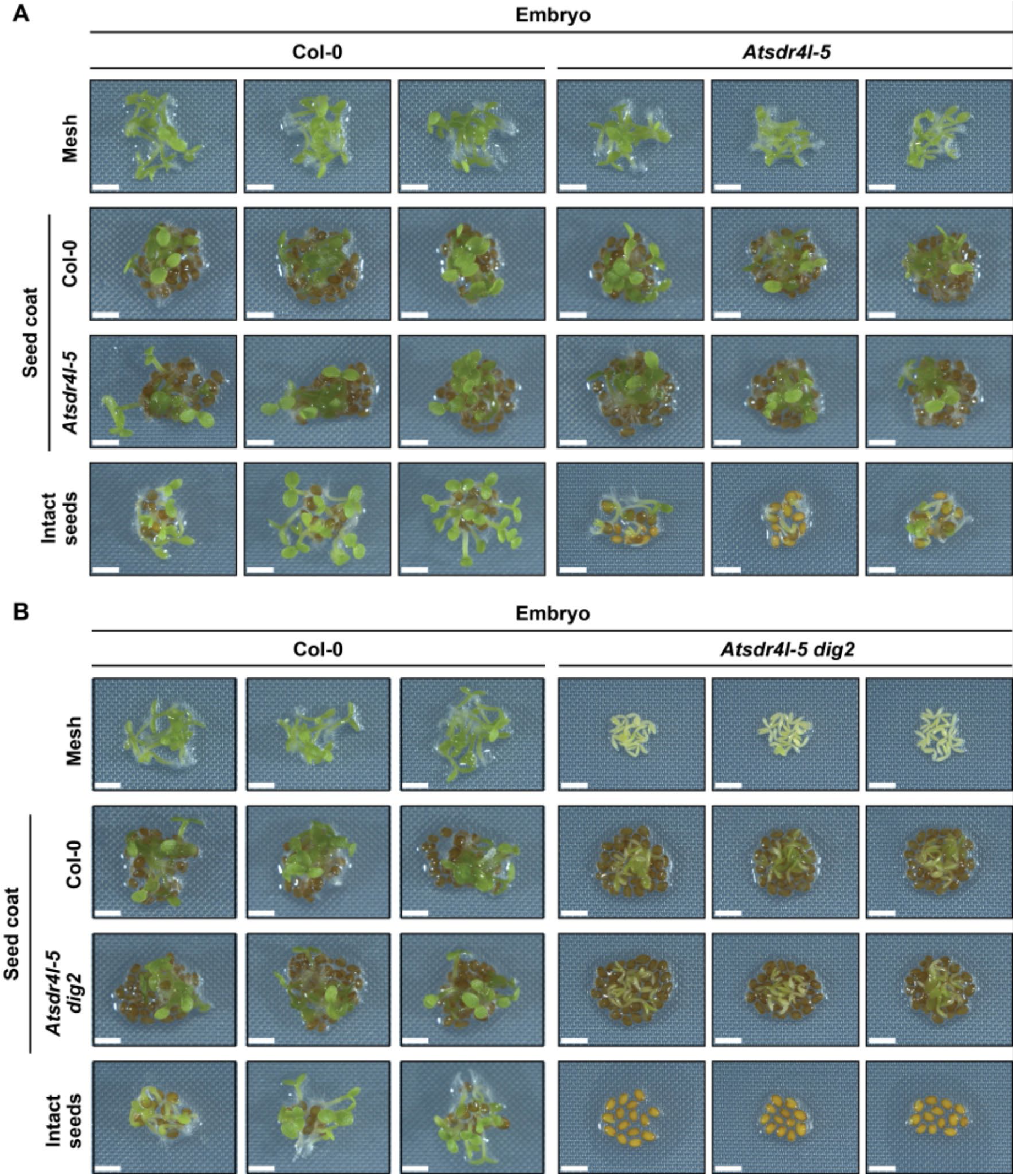
Seed germination arrest in *Atsdr4l-5 dig2* is contributed by both the embryo and the seed coat. (A,B) Establishment of Col-0 and *Atsdr4l-5* (A) or *Atsdr4l-5 dig2* (B) embryos directly on the medium (with mesh), and the seed coats of Col-0 and the corresponding mutant, and intact seeds at 3 DAI on 1XLS 0.7 % (w/v) phytoagar medium. Each column represents one set of biological replicates. Scale bar = 2 mm.

### *Atsdr4l dig2* embryo is more sensitive to osmotic stress and ABA

To examine whether *Atsdr4l-5 dig2* embryo has elevated sensitivity to ABA, we compared the growth of dissected wild-type and mutant embryos in the presence of 300 mM D-mannitol or 0.1 µM ABA (Fig. 3). On mock plates, approximately a third of *Atsdr4l-5 dig2* embryos lacked hairs at the collet zone between root and hypocotyl, contrastingly to the presence of collet hairs in all of Col-0, *Atsdr4l-5*, and *Atsdr4l-5 dig1* embryos. All mutant lines exhibited more severely arrested seedling establishment in response to osmotic stress or ABA. Additionally, fewer *Atsdr4l-5* and *Atsdr4l-5 dig1* embryos developed collet hairs than Col-0 while there was no collet hair growth in any *Atsdr4l-5 dig2* embryos. Thus, AtSDR4L synergistically works with its paralogs, in particular DIG2, to reduce embryo’s sensitivity to ABA and osmotic stress and facilitate the phase transition from seeds to seedlings. We also examined dissected embryos’ response to ABA analogues. Interestingly, whole seeds and embryos responded differently to the antagonists, and wild-type and *Atsdr4l-5 dig2* embryos exhibited ABA analogue-specific responses. Whereas seed germination and seedling growth of *Atsdr4l-5 dig2* were promoted by the ANT and ABA1019 (Fig. 1, S1B-C), growth of *Atsdr4l-5 dig2* embryos was mildly promoted by ANT and completely inhibited by ABA1019 (Fig. S3A-C). Growth of wild-type Col-0 embryos was modestly inhibited by ANT and completely inhibited by ABA1019, although Col-0 seedlings still established faster than *Atsdr4l-5 dig2* on mock medium and medium supplemented with 50 µM ANT. These data imply different ABA sensitivity in the endosperm and the embryo of wild type and *Atsdr4l-5 dig2* mutant, thereby resulting in tissue-and genotype-specific responses to a high level of ABA analogues.

**Fig. 3.**
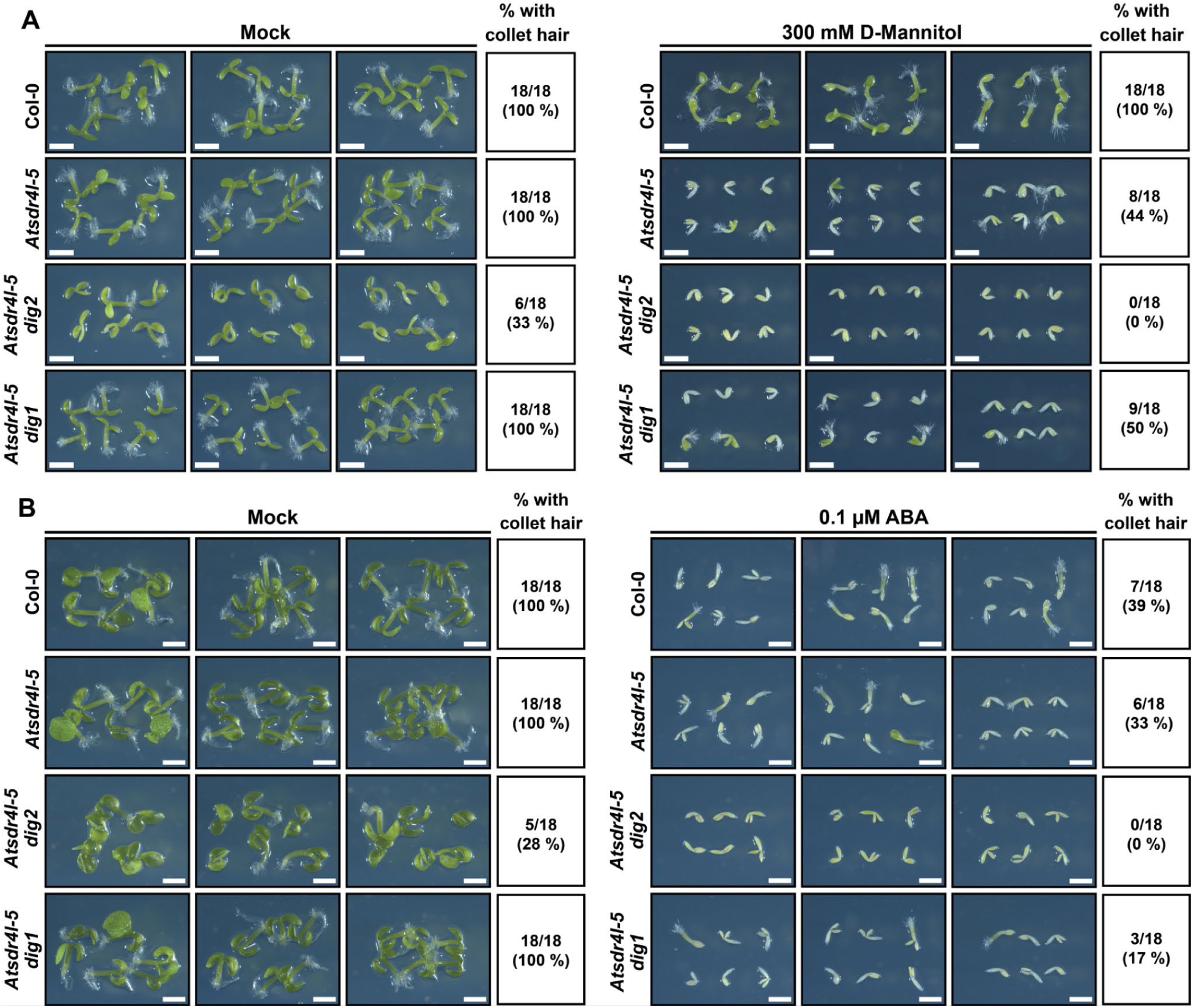
*Atsdr4l-5 dig2* embryos are hypersensitive to osmotic stress and ABA. (A) Embryos at 3 DAI on mock- (left) and 300 mM D-Mannitol- (right) treated 1X LS 0.7 % (w/v) phytoagar medium. (B) Embryos at 7 DAI on mock- (left) and 0.1 μM ABA- (right) treated 1X LS 0.7 % phytoagar (w/v) medium. Each column represents one set of biological replicates. Scale bar = 2 mm.

### *Atsdr4l dig2* embryo and seed coat show both similar and unique misregulation of genes, including *AGL* transcription factors

The results of seed coat bedding assay and hypersensitivity of dissected *Atsdr4l-5 dig2* embryo to ABA and osmotic stress prompted us to examine the transcriptomes of seed compartments in wild-type and the double mutant. We sequenced RNA-seq libraries prepared using dissected seed coats and embryos of Col-0 and *Atsdr4l-5 dig2*. Because the seed coat compartment contains 5 layers of dead testa cells and 1 layer of live endosperm cells, most if not all RNA is expected to be from the endosperm. Hierarchical clustering revealed that tissue type and genotype contribute to most differences between the samples (Fig. S4A). The endosperm marker *EARLY-PHYTOCHROME-RESPONSIVE1* (*EPR1*) is predominantly expressed in the seed coat (Fig. 4A), supporting clear separation of the tissues (Dubreucq et al., 2000). *AtSDR4L* is up-regulated in both mutant tissues, confirming previously reported self-repression (Wu et al., 2022; Lu et al., 2024). Several key regulators of seed maturation are de-repressed in the embryo and / or the seed coat of *Atsdr4l-5 dig2* (Fig. 4A). Notably, the basal expression level and the extent of up-regulation are locus- and compartment-specific. *ABI3* transcript is more abundant in the embryo, whereas *FUS3* transcript is more abundant in the seed coat of wild-type samples. The tissue preference of these two genes remains after being de-repressed in *Atsdr4l-5 dig2*. *MFT* and *DOG1* have comparable basal levels of expression between the two compartments, but *MFT* is only significantly up-regulated in *Atsdr4l-5 dig2* embryo, whereas *DOG1* is significantly up-regulated in both mutant tissues, with a stronger induction in the seed coat. The magnitude of differential expression is significantly and positively (Pearson correlation coefficient = 0.91, *p* < 2.2 x 10^-16^) correlated between the two tissues, with only a small set of genes oppositely differentially expressed (Fig. 4B). Consistent with delayed germination, genes related to positive regulation of germination arrest have significantly elevated expression in the mutant embryo and seed coat (Fig. 4B). The up-regulated genes in both tissues are enriched for biological processes related to the responses to hormonal, chemical and abiotic signals, with a subset of the genes showing embryo-specific functions (Fig. S4B). The down-regulated targets in both tissues are associated with responses to chemical and biotic stimuli, while those in embryo also have relations to cell wall modification, transcription and metabolic processes, and those in seed coat are linked with defense and toxin responses (Fig. S4B).

**Fig. 4.**
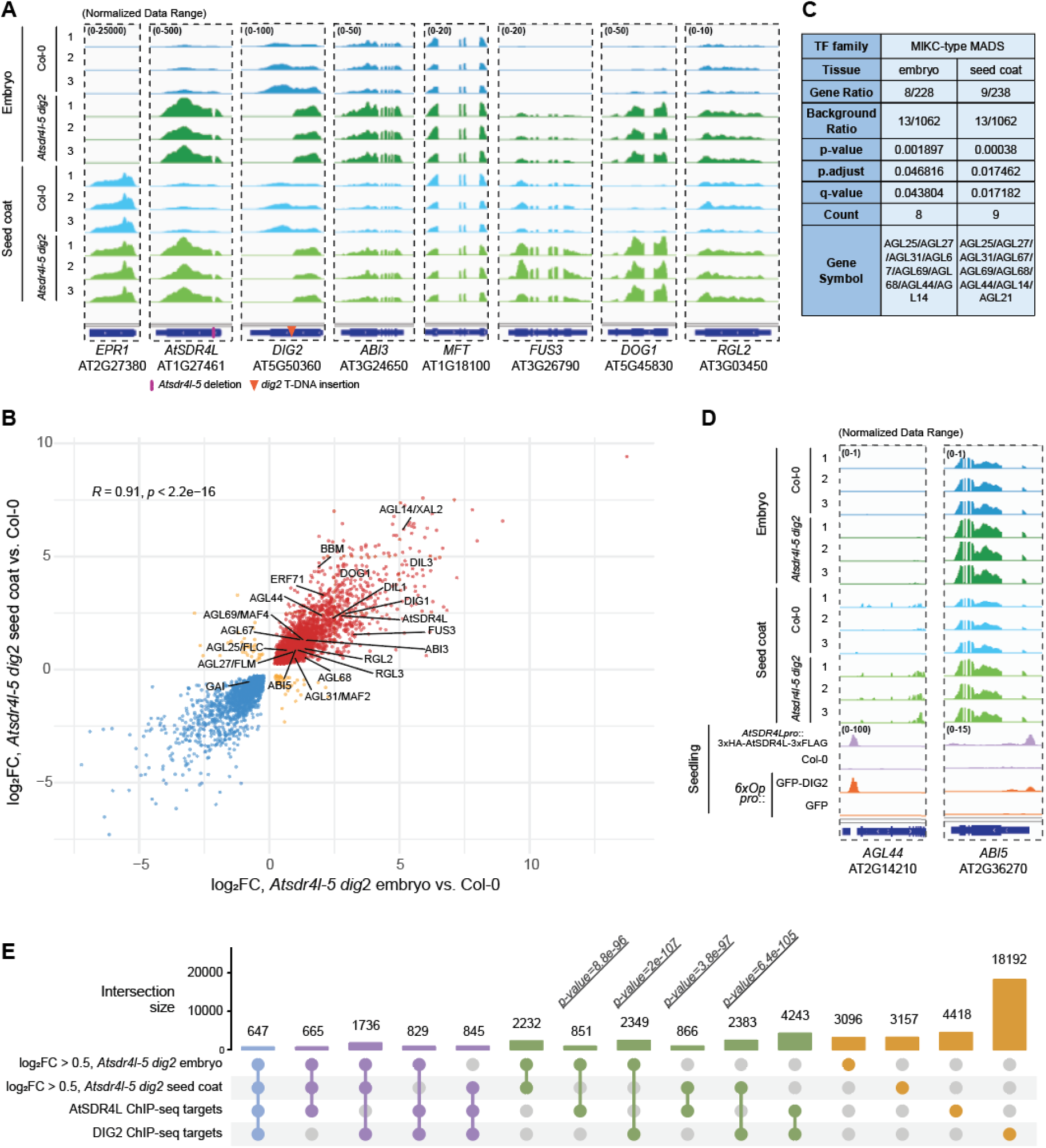
Mis-regulated genes in *Atsdr4l-5 dig2* embryo and seed coat include important regulators of germination. (A) Differential expression (DE) of *AtSDR4L*, *DIG2*, and selected regulators of germination in Col-0 and *Atsdr4l-5 dig2* embryo and seed coat shown in Integrative Genomics Viewer (IGV). Diamond and triangle labels mark the position of segmental deletion in *AtSDR4L* and T-DNA insertion in *DIG2*, respectively. Numbers in parentheses specify the range of normalized data. (B) Scatterplot showing the correlation in the magnitudes of misregulation (log_2_FC computed against Col-0) for significantly DE genes in the *Atsdr4l-5 dig2* mutant embryo and seed coat. Selected transcriptional regulators of dormancy and germination and up-regulated *AGAMOUS-LIKE* (*AGL*) genes are annotated. (C) Enrichment of MIKC-type MADS domain transcription factor family in the up-regulated genes (adj.P.Val < 0.05, log_2_FC > 0.5) in both mutant embryo and seed coat. (D) IGV screenshots for *AGL44* and *ABI5*, with the RNA-seq tracks showing their up-regulation in the mutant tissues, and ChIP-seq tracks showing the evidence of direct binding by AtSDR4L (SRR26666712) and DIG2 (SRR3418156) near the genes in young seedlings. Numbers in parentheses indicate the normalized data range. (E) UpSet plot for the intersection among the significantly up-regulated genes in *Atsdr4l-5 dig2* seed compartments and the genes with direct binding of AtSDR4L or DIG2 within 3-kb in the seedling ChIP-seq. P-values indicate the significance of overlap. Bars and dots are colour-coded according to the numbers of intersected groups (light blue = shared by all four groups, purple = shared by only three of the groups, green = shared by two groups, orange = total number of genes in each group alone).

To examine the changes in transcriptional regulation in *Atsdr4l-5 dig2* seeds, we obtained lists of TF families from AGRIS and PlantTFDB (Davuluri et al., 2003; Tian et al., 2019). A large number of TFs are misregulated in the seed compartments of *Atsdr4l-5 dig2*, suggesting extensive transcriptional rewiring. We identified MIKC-type MADS-box genes as the only group enriched in both the embryo and the seed coat (q-value < 0.05, hypergeometric test) among the up-regulated TFs (log_2_FC > 0.5, adj.P.val < 0.05) (Fig. 4B, 4C), and no TF family is enriched in the down-regulated TFs. Among 13 MIKC-type MADS-box TFs that are expressed in mature seeds, 8 are up-regulated in both seed compartments of *Atsdr4l-5 dig2*, including *AGL14*, *AGL25*, *AGL27*, *AGL31*, *AGL44*, *AGL67*, *AGL68*, and *AGL69.* Additionally, *AGL21* is specifically up-regulated in the seed coat of *Atsdr4l-5 dig2*. By intersecting the up-regulated TFs with previously generated ChIP-seq data of AtSDR4L and DIG2 in young seedlings (Song et al., 2016; Lu et al., 2024), we identified strong binding of both AtSDR4L (-log_10_[qValue] = 294, Benjamini-Hochberg correction) and DIG2 (-log_10_[qValue] = 1446) to the upstream of *AGL44* (Fig. 4D). Similarly, strong binding by both AtSDR4L (-log_10_[qValue] = 102) and DIG2 (- log_10_[qValue] = 88) was observed upstream of *ABI5* (Fig. 4D), suggesting direct de-repression contributes to the up-regulation of these two TFs in *Atsdr4l-5 dig2* seeds. Global intersection of differentially expressed genes in *Atsdr4l-5 dig2* seed compartments and binding sites of AtSDR4L and DIG2 further confirms that these co-repressors bind more frequently to genes up-regulated in the mutant, in contrast to those that are down-regulated (Fig. 4E, S4C), corroborating their roles in transcriptional repression.

### Multiple phytohormones are mis-regulated in Atsdr4l-5 dig2 seeds

Our transcriptome analysis showed that many genes regulating hormone metabolism are mis-expressed in the seed compartments of *Atsdr4l-5 dig2*. We focused on genes related to ABA, auxin, and ethylene because of their roles in dormancy and germination. Multiple *NCED* genes are up-regulated in both seed compartments of *Atsdr4l-5 dig2*, and their transcripts accumulate to a higher level in the embryo than in the seed coat (Fig. 5A, S5A). *CYP707A2* is only differentially expressed in the embryo of the double mutant (Fig. 5A), and its up-regulation may suggest a feedback to an elevated level of ABA in *Atsdr4l-5 dig2* seeds. In Arabidopsis, indole-3-acetic acid (IAA), the most common form of natural auxin, is synthesized via multiple tryptophan-dependent and independent pathways, and the conversion to IAA from indole-3-pyruvic acid, indole-3-acetamide, and indole-3-acetonitrile are catalyzed by YUCCA (YUC), AMIDASE 1 (AMI1), and NITRILASE (NIT) enzymes, respectively (Mano and Nemoto, 2012). Genes encoding these enzymes are generally up-regulated in *Atsdr4l-5 dig2*. For instance, *YUC*2 and *NIT1* exhibit increased expression in both seed compartments of the double mutant (Fig. 5A), *NIT2* is up-regulated only in the embryo (Fig. 5A), and *AMI1* is up-regulated in the embryo but down-regulated in the seed coat. The trend is more convoluted for the biosynthesis of 1-aminocyclopropane-1-carboxylic acid (ACC), the immediate precursor of ethylene due to the opposite changes in the transcript levels of *ACC SYNTHASE 2* (*ACS2*) and *ACS6* in *Atsdr4l-5 dig2* seeds (Fig 5A).

**Fig. 5.**
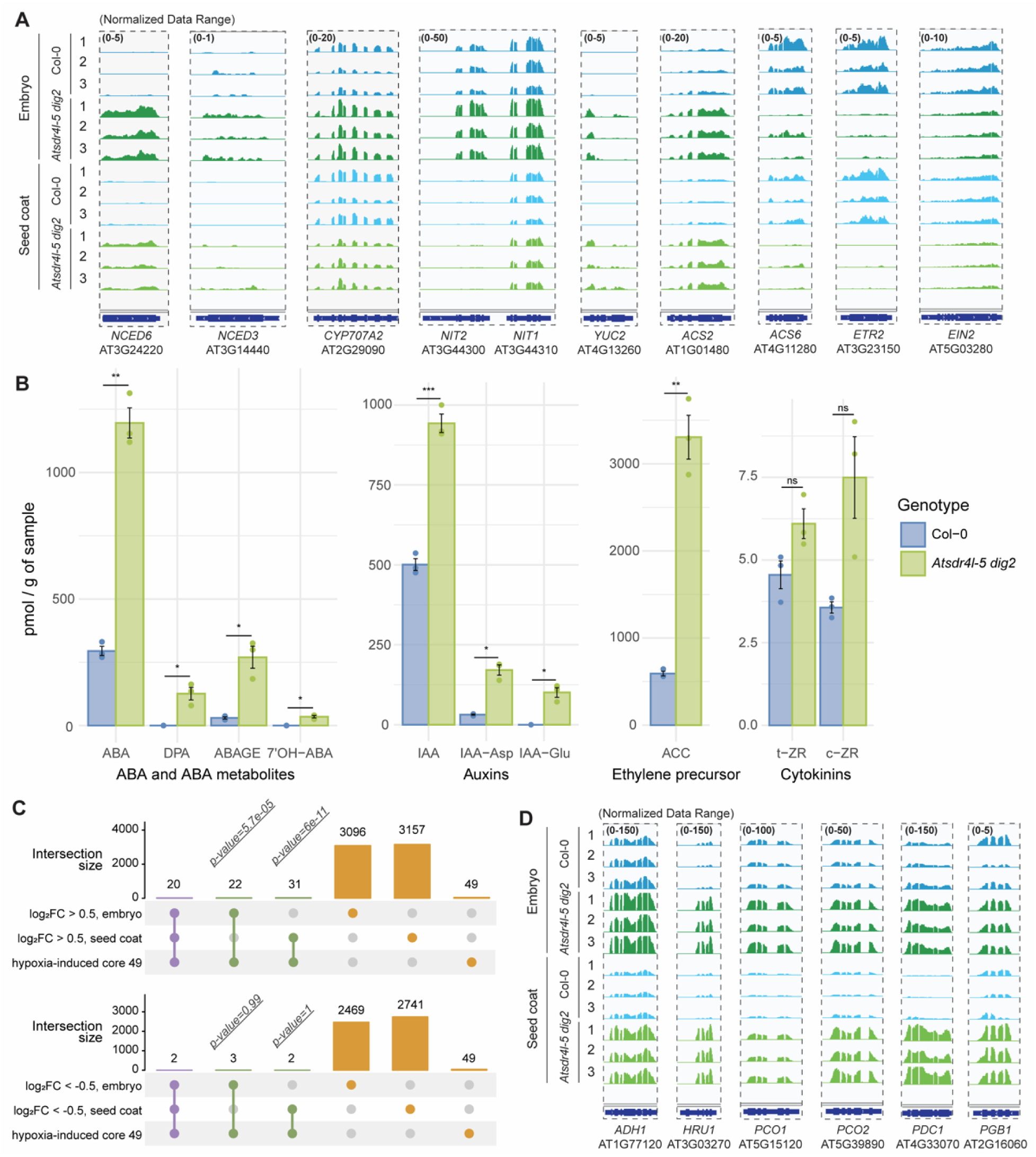
Multiple phytohormones and hypoxia-responsive genes are mis-regulated in *Atsdr4l-5 dig2* seeds. (A) Selected genes involved in ABA, auxin, and ethylene biosynthesis or catabolism, as well as factors in the ethylene signaling pathway are shown in Integrative Genomics Viewer (IGV). Numbers in parentheses indicate the normalized data range. (B) Bar plots for the quantification (pmol / g of sample) of ABA and ABA metabolites, auxin and its derivatives, ethylene precursor and cytokinins in seeds. Error bars represent mean ± SEM. \**P* < 0.05, \*\**P* < 0.01, \*\*\**P* < 0.001 (3 biological replicates, two-tailed Student’s t-test). (C) UpSet plots for the size of intersection among core 49 genes induced by hypoxia and the significantly up- and down-regulated genes in *Atsdr4l-5 dig2* embryo and seed coat. P-values indicate the significance of overlap. Bars and dots are colour-coded according to the numbers of intersected groups (purple = shared by all three groups, green = shared by two groups, orange = total number of genes in each group alone). (D) IGV screenshots for selected hypoxia-responsive genes that are significantly up-regulated in the *Atsdr4l-5 dig2* mutant seed tissues.

To confirm the effects of altered expression of hormone metabolism genes in *Atsdr4l-5 dig2*, a comprehensive hormone analysis was carried out using mature seeds of Col-0 and *Atsdr4l-5 dig2*. Previously, Zheng *et al*. showed that dry seeds of *Atsdr4l dig2* (known as *sfl1 sfl4* in their study) contain more ABA and auxin (Zheng et al., 2022). Consistently, we also observed that ABA and indole-3-acetic acid (IAA) levels in *Atsdr4l-5 dig2* seeds are 4.1 times and 1.9 times as much as those in wild-type Col-0 seeds, respectively (Fig. 5B). Elevated levels of ABA and auxin in *Atsdr4l-5 dig2* corroborate the elevated transcript abundance of *NCED* genes and the double mutant’s delayed germination. Consistent with the elevated level of ABA, expression of ABA signaling and ABA response genes in *Atsdr4l-5 dig2* seeds are largely similar to young Col-0 seedlings treated with exogenous ABA (Fig. S5B). Additionally, we found that dry seeds of *Atsdr4l-5 dig2* over-accumulate oxidized (7’-OH ABA, DPA) and conjugated (ABAGE, IAA-Asp, IAA-Glu) forms of ABA and auxin (Fig. 5B), suggesting that the deactivation pathways of ABA and auxin are up-regulated in *Atsdr4l-5 dig2* seeds. The over-accumulation of the oxidized forms of ABA is consistent with the up-regulation of *CYP707A2*.

Surprisingly, we observed a strong over-accumulation of the ethylene precursor ACC in *Atsdr4l-5 dig2* seeds (Fig. 5B). Ethylene or ACC is known to break dormancy in many species including *Arabidopsis thaliana* (Corbineau et al., 2014). We examined the expression of ethylene-responsive genes to infer the level of ethylene in the double mutant seeds. At the transcript level, ethylene induces the expression of its receptor *ETHYLENE RESPONSE 2* (*ETR2*) (Chen et al., 2007), a positive regulator of seed germination (Wilson et al., 2014). Transcript abundance of *ETR2*, along with signal transducer *ETHYLENE INSENSITIVE 2* (*EIN2*) and transcription factors *EIN3* and *EIN3-LIKE1* (*EIL1*) are all down-regulated in *Atsdr4l-5 dig2* seeds (Fig. 5A). To elucidate whether ethylene signaling is attenuated in *Atsdr4l-5 dig2*, We carried out germination assays on medium supplemented without and with 50 µM of ACC. Our results show that while 50 µM ACC treatment very modestly increases the germination of Col-0 seeds at 1 DAI, ACC has a larger effect in promoting the germination of *Atsdr4l-5 dig2* seeds (Fig. S5C). However, germination of *Atsdr4l dig2* seeds is not restored to the same level as wild-type Col-0 seeds even with a high level of exogenous ACC. These results suggest that the double mutant is still responsive to the ethylene precursor ACC, which is consistent with the known effect of ethylene in promoting seed germination through modulation of ABA (Guo et al., 2023). The reduction in the transcript level of ethylene signaling components likely contributes to the reduction in seed germination of the double mutant. The upregulation of ACC in *Atsdr4l dig2* is likely caused by a release of negative feedback due to repressed ethylene signaling (Thain et al., 2004). Hormone analysis also showed that the level of cis-zeatin riboside (c-ZR) in *Atsdr4l-5 dig2* is almost two times as high as in Col-0 seeds (Fig. 5A). Unlike trans-zeatin-type cytokinins, cis-zeatin and c-ZR are often considered less biologically active and are more abundant in growth-limiting tissues (Gajdošová et al., 2011; Schäfer et al., 2015). Although unlike the aforementioned hormones, the increase in c-ZR is statistically insignificant (p > 0.05, two-tailed Student’s t-test), the trend is consistent with the reduced growth potential of the double mutant. Collectively, these data showed that changes in transcript abundance of biosynthesis and signaling genes and over-accumulation of ABA and auxin support enhanced germination arrest in *Atsdr4l-5 dig2* seeds.

### Atsdr4l dig2 seeds exhibit elevated hypoxia responses

The elevated level of ACC and increased transcript abundance of *ACS2* prompted us to examine related changes such as hypoxia (Hartman et al., 2021). We observed differential expression of multiple *Group VII Ethylene Response Factor* (*ERF-VII*) genes that encode positive regulators of hypoxia responses and negative regulators of germination (Gibbs et al., 2015; Gasch et al., 2016). Because ERF-VII TFs are modified post-translationally (Giuntoli and Perata, 2018), we inferred their activities in *Atsdr4l-5 dig2* seeds by examining the expression of their downstream genes. We analyzed a list of “core 49” hypoxia-responsive genes (HRGs) that are induced by hypoxia in Arabidopsis root, shoot, and root tip (Mustroph et al., 2009), and observed that among 42 of the “core 49” genes that are expressed in seeds, a significant portion overlaps with the up-regulated genes in the embryo or seed coat of *Atsdr4l-5 dig2* (Fig. 5D). By contrast, only a few genes such as *ETR2* are down-regulated in *Atsdr4l-5 dig2*, plausibly resulting from the mis-regulation of ethylene signaling in the mutant seeds. Examination of the browser tracks further confirmed that several well-characterized HRGs are up-regulated in both compartments of *Atsdr4l-5 dig2* seeds, including *ALCOHOL DEHYDROGENASE 1* (*ADH1*), *HYPOXIA RESPONSIVE UNIVERSAL STRESS PROTEIN 1* (*HRU1*), *PLANT CYSTEINE OXIDASE 1* (*PCO1*), *PCO*2, *PYRUVATE DECARBOXYLASE 1* (*PDC1*), and *PHYTOGLOBIN 1* (*PGB1*) (Fig. 5E). Additionally, we examined by RT-qPCR the expression of *PCO1* and *PGB1* levels in dry seeds, and showed these genes are up-regulated in *Atsdr4l-5 dig2* prior to imbibition (Fig. S5D). These data support increased hypoxia or elevated hypoxia responses in *Atsdr4l-5 dig2* seeds.

## Discussion

### Hormonal changes and a possible increase in hypoxia in seed compartments promote germination arrest of *Atsdr4l dig2*

In this study, we utilized a combination of assays to examine how *Atsdr4l dig2* mutant acquired strong seed germination arrest. The difference in the severity of germination delay between *Atsdr4l dig2* and *Atsdr4l dig1* is consistent with a higher transcript abundance of *DIG2* than *DIG1* towards late seed maturation compared to the rest of the family members (Gazzarrini and Song, 2024), implying that the paralogs in the DIG subclade of this gene family might be subfunctionalized. Arabidopsis seeds have non-deep physiological dormancy, and the balance between dormancy and germination is fine-tuned by phytohormones (Finch-Savage and Leubner-Metzger, 2006; Penfield, 2017). The ABA biosynthetic genes *NCED6* and *NCED3* and ABA-responsive regulators such as *ABI3*, *ABI5*, and *DOG1* are up-regulated in both seed compartments of *Atsdr4l-5 dig2*, and *ABI3* and *MFT* transcripts accumulate to a higher abundance in the embryo than the seed coat (Fig. 4A). The transcriptome changes corroborate elevated levels of ABA and its oxidized or conjugated metabolites (Fig. 5A-B). The higher abundance of the oxidized ABA metabolites (7’-OH ABA, DPA) in *Atsdr4l-5 dig2* seeds may be explained by the up-regulation of the catabolic gene *CYP707A2* in the embryo (Fig. 5A) (Nambara and Marion-Poll, 2005; Nambara et al., 2010; Bai et al., 2022). ABA antagonist treatments further confirmed that ABA is the major determinant for inhibited germination and subsequent seedling establishment in *Atsdr4l-5 dig2*. The defects in germination and axis elongation of intact *Atsdr4l-5 dig2* seeds are alleviated by antabactin and ABA1019 (Fig. 1, S1B). Contrastingly, ABA1019 completely suppressed the growth of dissected Col-0 and mutant embryos (Fig. S3). Such differences in ABA1019’s effects on intact seeds and dissected embryos could be dose- or tissue-dependent, and ABA1019 was reported to function as both an antagonist and agonist (Diddi et al., 2021). Previous studies reported the opposing effects from the ligand-receptor interaction on pyrabactin as both an ABA antagonist and agonist (Peterson et al., 2010; Yang et al., 2020a). As such, we speculate that the opposite effect of ABA1019 on whole seeds and dissected embryos might be contributed by the receptor-specific mode of actions and by the differences in the expression of various ABA receptors in seed coat and the embryo (Gonzalez-Guzman et al., 2012). This speculation is also supported by the compartment-specific differential expression of a subset of ABA receptor transcripts such as *PYL1* and *PYL12* (Fig. S5B). Collectively, gene expression profiling supports the increased responsiveness to ABA in *Atsdr4l-5 dig2* seeds. Preferential up-regulation of many positive regulators of ABA signaling is consistent with the seed coat bedding assay results that *Atsdr4l-5 dig2* embryo is dormant even in the absence of the seed coat (Fig. 2, S2).

Changes in other hormones such as auxin are also supported by the transcriptome data. The significantly elevated expression of *YUC2* and *NIT1* in *Atsdr4l-5 dig2* (Fig. 5A) may be responsible for the overaccumulation of IAA and its amino acid conjugates (Fig. 5B). Additionally, *NIT2* and *AMI1* are up-regulated in *Atsdr4l-5 dig2* embryo, suggesting that multiple pathways are activated in the mutant embryo to increase IAA biosynthesis. Therefore, impeded seed-to-seedling transition in *Atsdr4l-5 dig2* likely results from elevated auxin level and the interplay between auxin and ABA to enhance germination arrest and inhibit the elongation of the embryonic axis (Belin et al., 2009; Liu et al., 2013). Consistent with delayed germination of the double mutant, several expansin genes (Weitbrecht et al., 2011; Stamm et al., 2012), including *EXPANSIN3* (*EXP3*), *EXP8*, *EXP20*, are down-regulated in *Atsdr4l-5 dig2* seeds. *EXP8* is weakly expressed mainly in the seed coat of Col-0, possibly activated by the brief imbibition during seed dissection, whereas its expression dropped noticeably in the seed coat of *Atsdr4l-5 dig2* (Log_2_FC = - 3.83, p.adj = 0.0067, Benjamini-Hochberg correction) (Fig. 6A). Similarly, a significantly lower expression of *EXP3* was found in *Atsdr4l-5 dig2* embryo compared to that of Col-0 (Log_2_FC = - 2.54, p.adj = 0.0006). These results suggest hindered cell wall loosening in both seed compartments of *Atsdr4l-5 dig2*.

**Fig 6.**
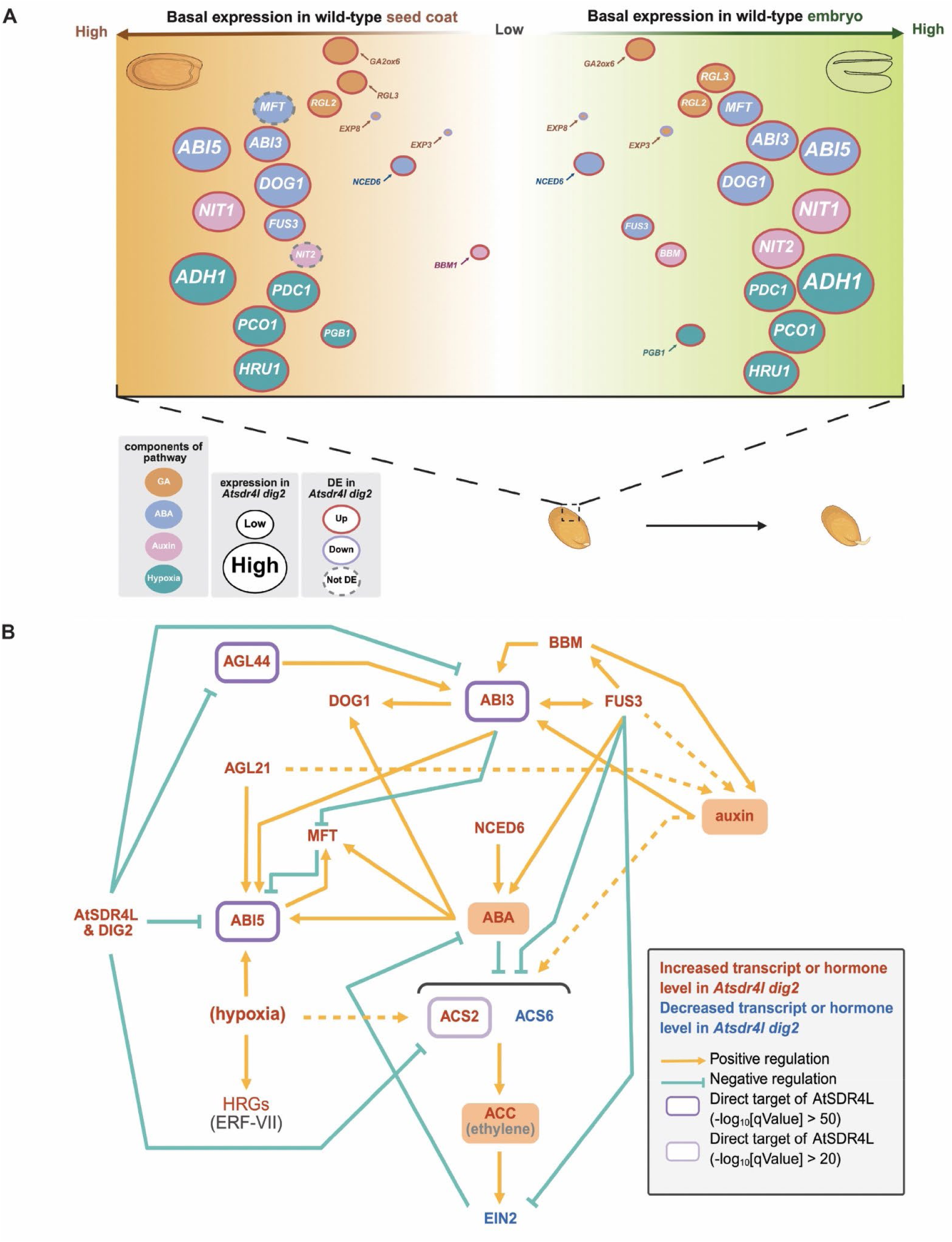
Proposed model for the tissue-specific regulation by AtSDR4L and DIG2 on selected dormancy, germination, and development genes in the embryo and seed coat. (A) In mature wild-type seeds, AtSDR4L works synergistically with DIG2 to repress the positive regulators of ABA and auxin biosynthesis and signaling, as well as the negative regulators in the GA signaling to promote germination and subsequent seedling establishment. The basal level of gene expression in the wild-type seed compartments, approximated by log_2_(FPKM), is indicated by the position of the gene along the horizontal axis. The size of the ellipses reflects transcript abundance of indicated genes in *Atsdr4l-5 dig2*, and ellipse outline indicates whether the gene is differentially expressed. Many of these genes exhibit preferential expression in either the seed coat or the embryo. Ellipse color reflects whether the gene is involved in biosynthesis, signaling, or downstream responses of indicated hormones. Genes related to multiple hormones are assigned to one. (B) AtSDR4L and DIG2 regulate the network of seed development and germination. Increased or decreased levels of transcripts or hormones in *Atsdr4l dig2* are marked in dark orange and blue colours, respectively. Positive and negative regulations among transcriptional transcriptional regulators and hormones based on literature are shown in yellow and teal colours, respectively. Solid arrows are derived from evidence specific to seed and dashed arrows are based on evidence in other tissues. Direct binding of AtSDR4L and DIG2 to the upstream region of their targets is shown in dark purple for -log_10_[qValue] higher than 50 and light purple for -log_10_[qValue higher than 20 of the AtSDR4L peaks.

Besides hormones, germination can also be regulated by hypoxia. Internal oxygen level is low in seeds (Borisjuk and Rolletschek, 2009), and increased hypoxia by physical barrier or metabolic changes may enhance germination arrest (Chandler et al., 2024; Pucciariello and Perata, 2024; Rolletschek et al., 2024). We observed up-regulation of a large number of HRGs, often in both seed compartments of *Atsdr4l-5 dig2*, suggesting activation of ERF VII TFs and increased hypoxia. This observation is consistent with the better germination of dissected embryos of *Atsdr4l-5 dig2* than intact mutant seeds (Fig. 2, S2), as the germination of isolated embryos are less sensitive to hypoxia in multiple species (Corbineau, 2022). Additionally, ERF VII TFs may activate *ABI5* to inhibit germination through crosstalk with ABA signaling (Gibbs et al., 2014). Seed’s sensitivity to hypoxia varies by species (Corbineau, 2022). Germination and coleoptile elongation may be stimulated by oxygen deprivation in certain rice cultivars (Magneschi and Perata, 2009). Therefore, it would be very interesting to examine the regulation of hypoxia response by AtSDR4L and its paralogs in a broad range of species.

### A transcriptional and hormonal network regulating seed-to-seedling transition

Pleiotropic phenotype reflects complex regulation, which is very common in the mutants of transcriptional regulators because of the large number of affected target genes and pathways. Therefore, comprehensive characterization of mutants will help dissect the pleiotropic phenotype and uncover complex regulatory networks. Our characterization of ethylene signaling in the *Atsdr4l-5 dig2* seeds provides an example of the collision of multiple hormonal and transcriptional regulation. Ethylene breaks dormancy and promotes germination through reducing ABA biosynthesis or sensitivity (Beaudoin et al., 2000; Ghassemian et al., 2000; Guo et al., 2023). Additionally, ethylene represses its receptor ETR1 and subsequently de-represses *ERF12*, the encoded TF of which recruits transcriptional co-repressor TOPLESS (TPL) to repress *DOG1* (Li et al., 2019). Hormone profiling reveals an increased level of ethylene-precursor ACC in the mutant seeds (Fig. 5B). This agrees with the up-regulation of *ACS2* in the embryo and seed coat of *Atsdr4l-5 dig2*, which could result from direct de-repression as binding of AtSDR4L (-log10[qValue] = 23) and DIG2 (-log10[qValue] = 827) was detected upstream of *ACS2*. Increased ACC in *Atsdr4l-5 dig2* may also indirectly result from auxin-dependent induction (Tsuchisaka and Theologis, 2004; Yang and Hoffman). Feedback regulation from reduced ethylene signaling will also increase ethylene biosynthesis (Thain et al., 2004). FUS3 negatively regulates ethylene signaling (Lumba et al., 2012) (Fig. 6B), and its up-regulation in *Atsdr4l-5 dig2* may down-regulate core ethylene signaling and responsive genes in the mutant seed compartments, including *ETR2*, *EIN2*, *EIN3*, and *EIL1* (Fig. 4A). However, exogenous ACC treatment showed that elevated ethylene biosynthesis is not sufficient to promote the germination of *Atsdr4l-5 dig2* seeds to the same level as wild-type Col-0 (Fig. S5C). This results indicate that the effect of elevated ACC must be limited by the down-regulation ethylene signaling genes, which may also enhance dormancy by alleviating ABA biosynthesis and responses (Beaudoin et al., 2000; Ghassemian et al., 2000; Guo et al., 2023). These data showed complex changes in ethylene biosynthesis and signaling to control dormancy in *Atsdr4l dig2*.

The direct targets of AtSDR4L and DIG2 provide additional examples of complex regulation (Fig. 4D). Previously, we reported that AtSDR4L and DIGs bind to the upstream regions of *ABI3* (Lu et al., 2024). Because the first three DAI is a critical developmental window for the seed-to-seedling transition in Arabidopsis (Lopez-Molina et al., 2001), the RNA-seq data generated using 4-day-old Col-0 and mutant seedlings (Wu et al., 2022) are informative to identify regulators of post-germinative development, but might miss important candidate genes that regulate germination. In this study, we intersected the binding data, especially the AtSDR4L binding data in 1-day-old seedlings that still retain many seed characteristics (Lu et al., 2024), with the differentially expressed genes in the seed compartments of *Atsdr4l-5 dig2*. We identified *AGL44* and *ABI5* as two additional promising targets of AtSDR4L and DIG2. *AGL44* encodes an MIKC-type MADS-box TF and negative regulator of seed germination in the presence of ABA, salt, or osmotic stress by modulating *ABI3* through direct targeting (Lin et al., 2020). *AGL21* is another MADS-box TF gene that is up-regulated in *Atsdr4l-5 dig2*, and AGL21 inhibits germination by activating *ABI5* (Yu et al., 2017). Collective knowledge from this study and from the literature show that AtSDR4L and DIG2 promote germination in Arabidopsis by repressing multiple negative regulators of germination, including both *AGL44* and downstream TFs of AGLs such as *ABI3* and *ABI5* (Fig. 6B). Similar coordinated regulation may help to repress embryonic programs. In embryonic tissues, BBM transactivates *ABI3* (Horstman et al., 2017), FUS3 targets *BBM* (Wang and Perry, 2013), and ABI3 and FUS3 target each other (Tian et al., 2020). Therefore, these three TFs may form a cross-activation loop (Fig. 6B). Additionally, auxin biosynthesis is required for somatic embryogenesis and is activated by BBM (Li et al., 2022). AGL21 and FUS3 have also been linked to auxin biosynthesis in root tissues (Tang et al., 2017; Yu et al., 2017). Therefore, AtSDR4L and DIG2 may restrain both the accumulation of auxin and the embryonic programs during the seed-to-seedling transition by repressing these transcriptional regulators directly or indirectly.

Taken together, we provide new insights into how AtSDR4L and its paralog DIG2 holistically coordinate hormonal, transcriptional, and hypoxic responses in seeds. AtSD4L and its paralogs may simultaneously repress multiple activators of seed dormancy, thus establishing a robust mechanism to break or attenuate the positive feedback among these regulatory nodes and promote seed-to-seedling transition. Homologs of AtSDR4L and DIG2 are broadly present in green lineages, and SDR4 orthologs are known to prevent preharvest sprouting in rice and wheat, two of the top three major staple crops (Sugimoto et al., 2010; Zhang et al., 2014; Chen et al., 2021). The functional contrast of SDR4 orthologs between monocot crops and the eudicot model species Arabidopsis underscores the the importance of understanding the regulatory mechanism of these genes in order to fulfill the high application potential of this gene family and their natural and engineered alleles for the optimization of desirable level of seed dormancy. The reported regulatory network may serve as a comparison basis to understand the regulation of germination and seedling establishment in various species, and potentially enrich the resource pools in breeding programs to engineer the seed qualities and seedling resilience under adverse conditions.

## Acknowledgement

The authors thank Sean Cutler (UCR) and Sue Abrams (USask) for the ABA analogues and discussions.The authors thank Hong Qiao (UT Austin) for discussions on ethylene and Travis Lee (Salk) for discussions on hypoxia. Hormone measurement was carried out by the National Research Council Canada. The model in Fig. 6 was prepared using BioRender. The authors thank funding support from the University of British Columbia Four Year Doctoral Fellowships (UBC 4YF) and NSERC Postgraduate Scholarship - Doctoral to B.L., UBC 4YF to D.G., UBC Work Learn International Undergraduate Research Awards to J.S., and NSERC RGPIN-2019-05039 and CFI JELF/BCKDF 38187 to L.S.

## Competing interests

The authors declare no conflict of interest in this study.

## Author contributions

L.S., B.L. and D.G. conceived and designed the experiments. B.L., D.G., and J.S. performed the experiments. B.L., D.G., and L.S. analyzed the data. B.L., D.G., and L.S. wrote the manuscript.

## SI Figures

**Fig. S1.**
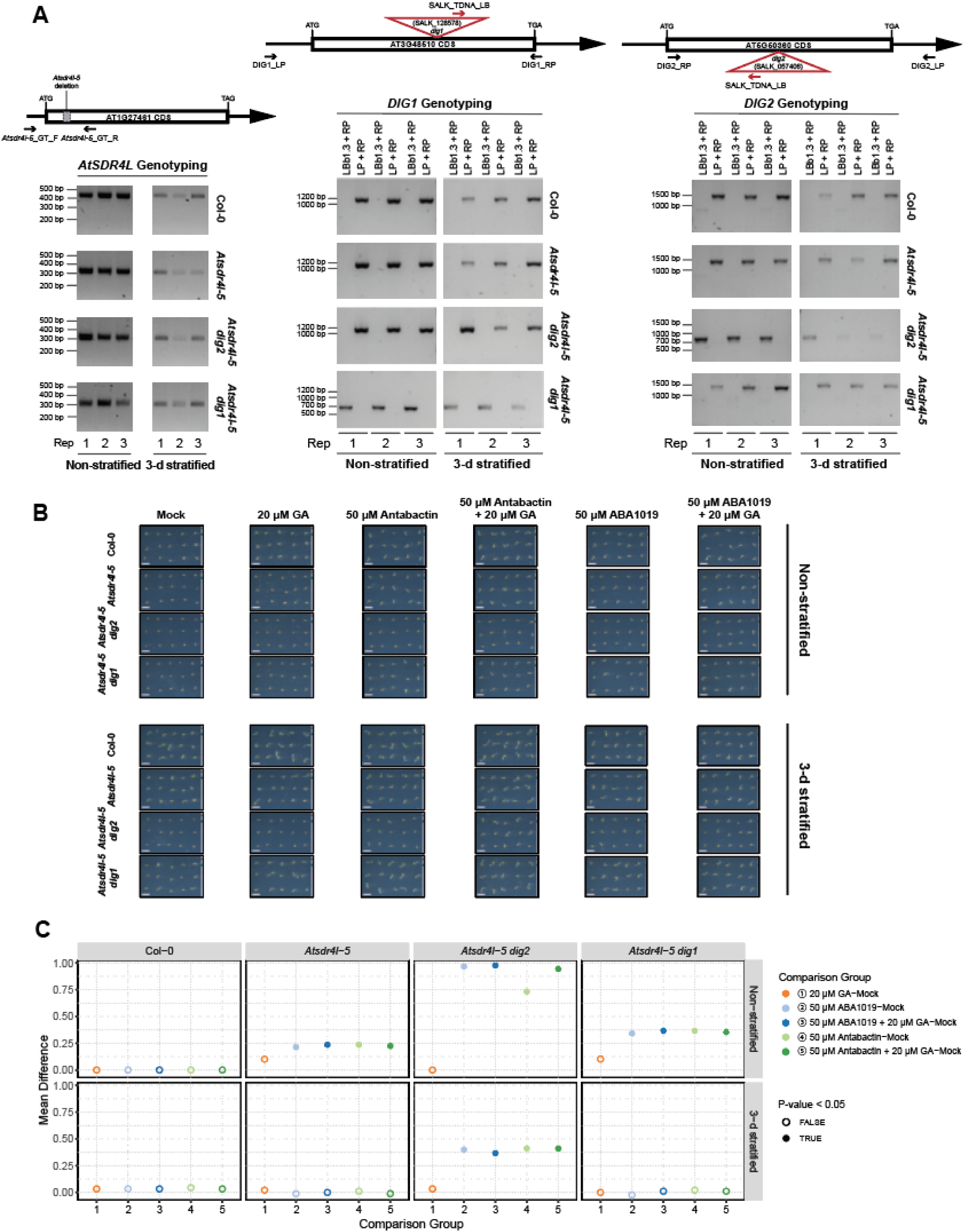
Seed germination arrest of *Atsdr4l-5 dig2* is alleviated by ABA antagonists. (A) A diagram of CRISPR/Cas9-mediated deletion that is harboured in *Atsdr4l-5* allele and T-DNA insertion map of *dig1* (SALK_128578) and *dig2* (SALK_057406) (top). Genotyping of Col-0, *Atsdr4l-5*, *Atsdr4l-5 dig2*, *Atsdr4l-5 dig1* used in this study (bottom). (B) Representative images of germination and embryonic axis growth of non-stratified (top) and 3-day-stratified (bottom) Col-0, *Atsdr4l-5*, *Atsdr4l-5 dig2*, *Atsdr4l-5 dig1* seeds and seedlings at 2 days after imbibition (2 DAI) post 2 weeks of after-ripening on 1xLS 0.7 % (w/v) phytoagar medium with treatments of 20 μM GA, 50 μM Antabactin, 50 μM Antabactin + 20 μM GA, 50 μM ABA1019, 50 μM ABA1019 + 20 μM GA-treated 1xLS 0.7 % (w/v) phytoagar medium. Each column represents one set of biological replicates. Scale bar = 4 mm. (C) ANOVA and Tukey’s HSD test results showing the difference in mean germination percentage between mock and each of the hormone/antagonist treatments for each genotype at 2 DAI. The solid and empty fills of the dots correspond to the statistical significance at *p* < 0.05.

**Fig. S2.**
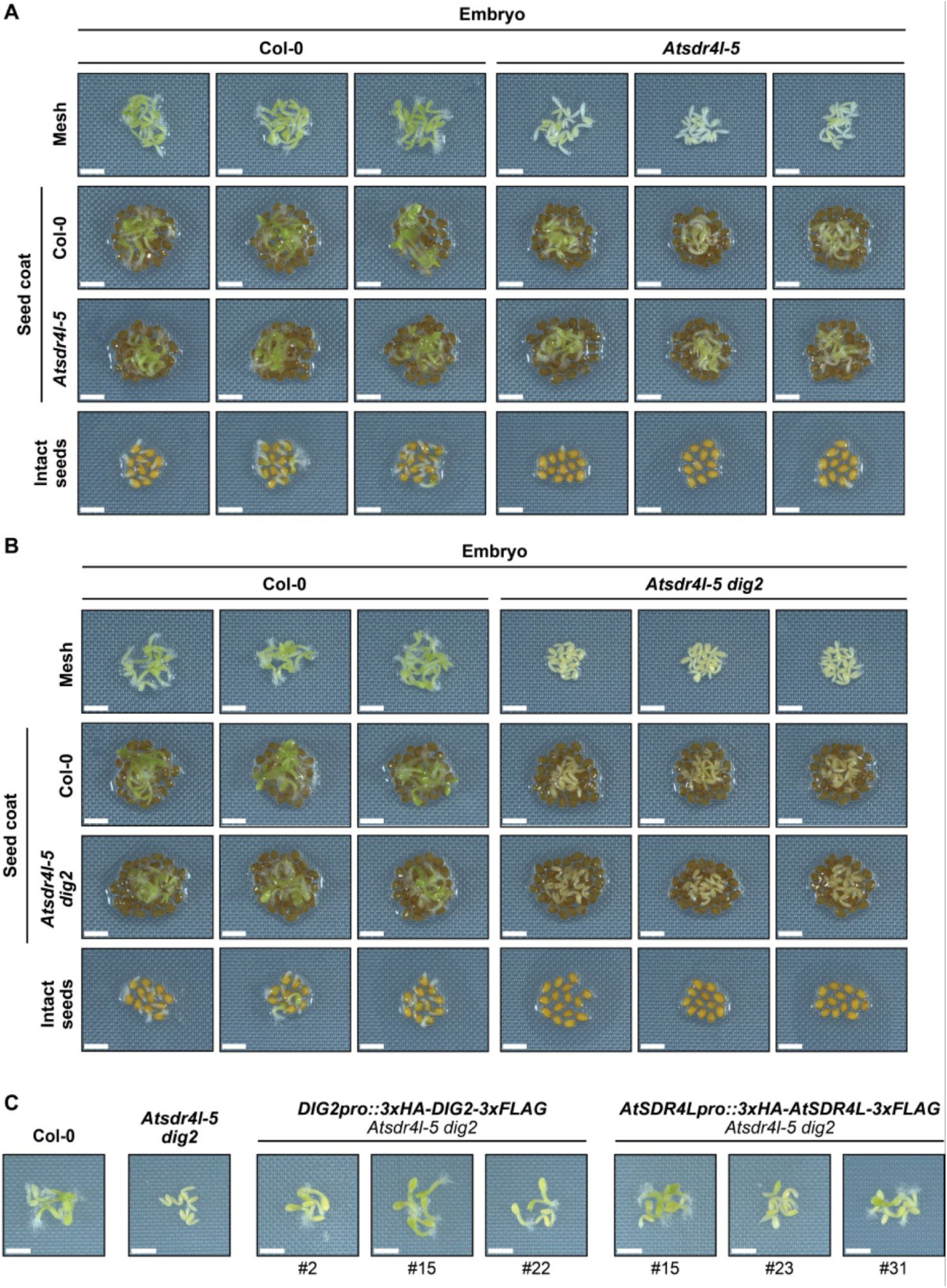
Inhibition of germination and seedling growth of *Atsdr4l-5 dig2* are exerted by the embryo and the seed coat. (A,B) Establishment of Col-0 and *Atsdr4l-5* (A) or *Atsdr4l-5 dig2* (B) embryos directly on the medium (with mesh), and the seed coats of Col-0 and the corresponding mutant, and intact seeds at 2 DAI on 1XLS 0.7 % (w/v) phytoagar medium. Each column represents one set of biological replicates. Scale bar = 2 mm. (C) Complementation of *Atsdr4l-5 dig2* mutant embryos from non-stratified T2 seeds by the vectors of *DIG2pro::3xHA-DIG2-3XFLAG* or *AtSDR4Lpro::3xHA-AtSDR4L-3XFLAG*. Embryos were imaged at 3 DAI. Scale bar = 2 mm.

**Fig. S3.**
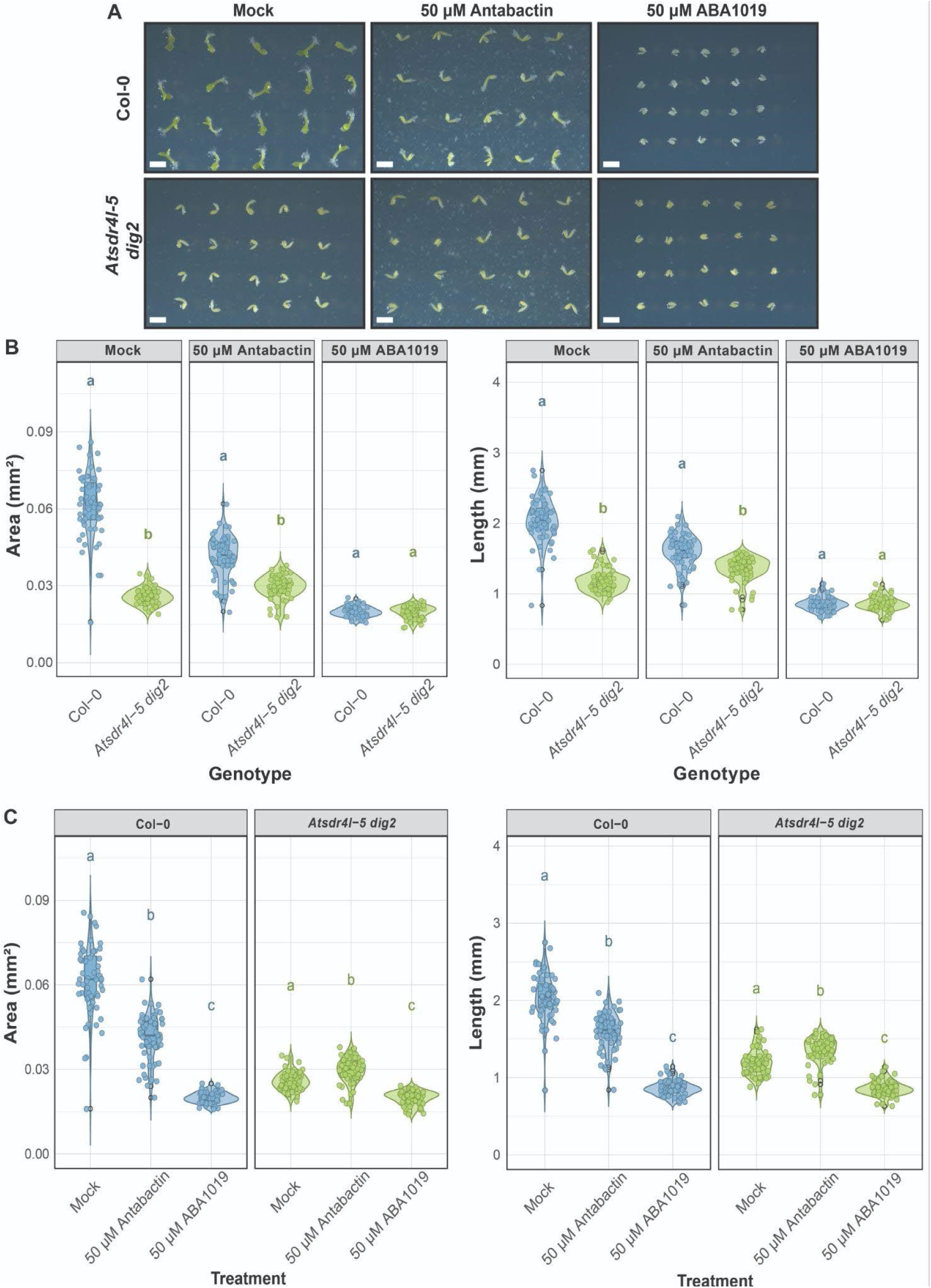
Wild-type and *Atsdr4l-5 dig2* embryos exhibit ABA analogue-specific responses in seedling establishment. (A) Representative images of embryos at 2 DAI on 1X LS 0.7 % (w/v) phytoagar medium with treatments of mock, 50 μM Antabactin, 50 μM ABA1019 after 2.5 weeks of after-ripening. Each column represents one set of biological replicates. Scale bar = 2 mm. (B - C) Total area (left) and embryonic axis length (right) of Col-0 and *Atsdr4l-5 dig2* seeds and seedlings statistically tested for the effect of genotype (B) and treatment (C). The black dots represent outliers. Different letters (*p* < 0.05) represent significant differences (n=20, 3 biological replicates, Kruskal-Wallis test, Dunn’s test).

**Fig. S4.**
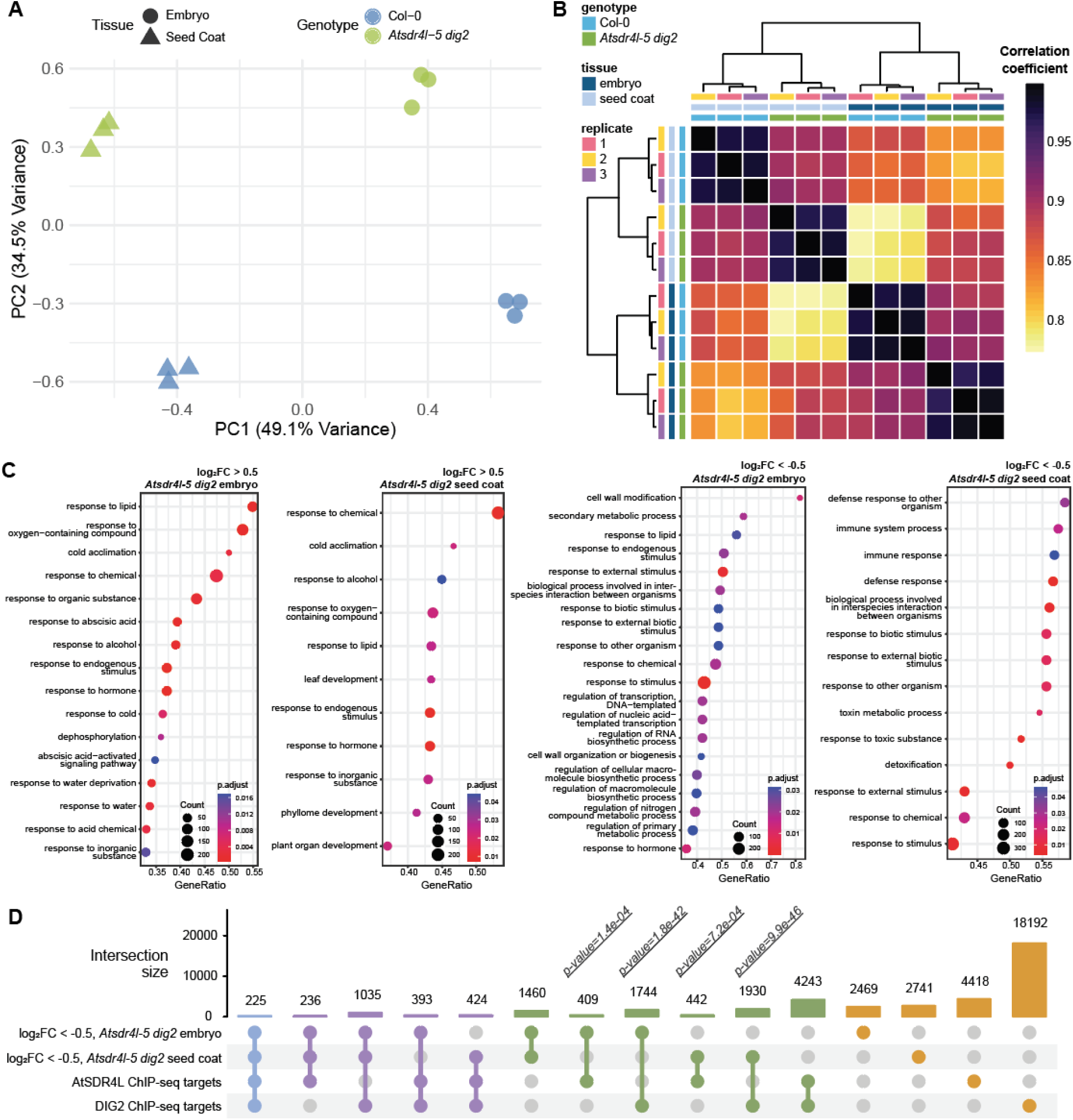
*Atsdr4l-5 dig2* transcriptome changes correlate with tissues and have distinct and similar functional categories. (A) Principal component analysis (PCA) by multidimensional scaling on the wild-type and mutant tissue-specific RNA-seq reads normalized by TMM (Trimmed Means of M-values) method. The shapes and colours of dots correspond to the tissue type and genotype. (B) Heatmap for the pairwise correlation among the Col-0 and *Atsdr4l-5 dig2* embryo and seed coat samples. Correlation matrix was computed using log_2_ transformed (with pseudo-count of 1) counts per million (CPM) values after filtering out genes with low read counts and normalizing library size by “Trimmed Means of M-values (TMM)” method. Colour scale represents the correlation coefficients. (C) Dot plots with the results of gene set enrichment analysis (GSEA) for Gene Ontology (GO) terms (biological process) associated with significantly up-regulated (adj.P.Val < 0.05, log_2_FC > 0.5) and down-regulated (adj.P.Val < 0.05, log_2_FC < -0.5) genes in *Atsdr4l-5 dig2* embryo and seed coat. The size of the dots is proportional to the number of genes with the corresponding GO annotation, and the colour represents the scale of adjusted p-values. (D) UpSet plot for the size of intersection among the significantly down-regulated genes in *Atsdr4l-5 dig2* seed compartments and the genes with direct binding of AtSDR4L or DIG2 within 3-kb in the seedling ChIP-seq. P-values indicate the significance of overlap. Bars and dots are colour-coded according to the numbers of intersected groups (light blue = shared by all four groups, purple = shared by only three of the groups, green = shared by two groups, orange = total number of genes in each group alone).

**Fig. S5.**
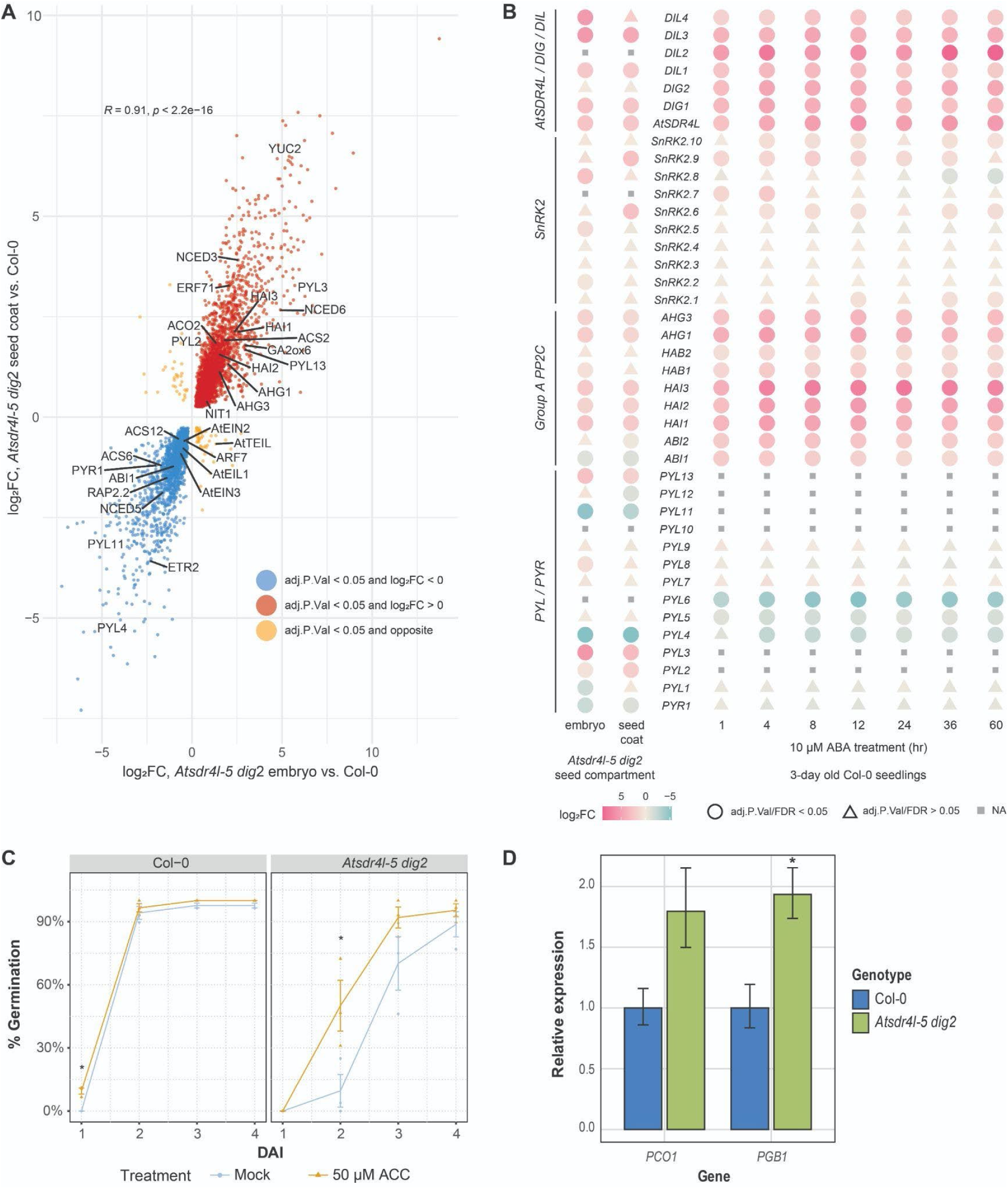
*Atsdr4l-5 dig2* seed compartments exhibit both shared and tissue-specific transcriptome changes for hormone-related genes. (A) Scatterplot with the log_2_FC values between mutant and Col-0 for significantly DE genes in the *Atsdr4l-5 dig2* mutant embryo and seed coat. Colour of the dots are labelled for the directions of misregulation. Selected genes involved in regulating ABA, GA, auxin, and ethylene biosynthetic and signaling pathways are annotated. (B) Dotted heat scale reflecting the up-regulation or down-regulation of genes encoding components of ABA signaling pathways in *Atsdr4l-5 dig2* embryo and seed coat, as well as in ABA-treated Col-0 seedlings (data from GSE80565). *PYR*/*PYL*: *PYRABACTIN RESISTANCE1*/*PYR1-LIKE* ABA receptors; *PP2C*: protein phosphatase 2C; *SnRK2*: *SNF1-RELATED PROTEIN KINASE2*; *AtS*DR4L/DIG/DIL: *ARABIDOPSIS THALIANA SEED DORMANCY 4-LIKE*/*Dynamic Influencer of Gene Expression*/*DIG-Like*. Shapes indicate whether the differential expression is significant in the corresponding tissue or treatment. Genes without detectable expression are indicated by NA. (C) Percentage of seed germination over days after imbibition (DAI) of non-stratified and 3-day cold stratified Col-0, *Atsdr4l-5*, *Atsdr4l-5 dig2* and *Atsdr4l-5 dig1* seeds on 1X LS 0.7 % (w/v) phytoagar medium with treatments of mock and 50 μM ACC. Error bars represent mean ± SEM (26 ≤ n ≤ 30, 3 biological replicates; \**p* < 0.05, one-tailed Student’s t-test). (D) *PCO1* and *PGB1* mRNA levels in ∼ 20 DAP seeds of Col-0 and *Atsdr4l-5 dig2*. *PCO1* and *PGB1* transcript accumulation was normalized to *ACTIN8*. Error bar represents a standard error of the mean (SEM) from 3 biological replicates (\**p* < 0.05, Student’s t-test).

## SI Tables

**Supplementary Table 1:**
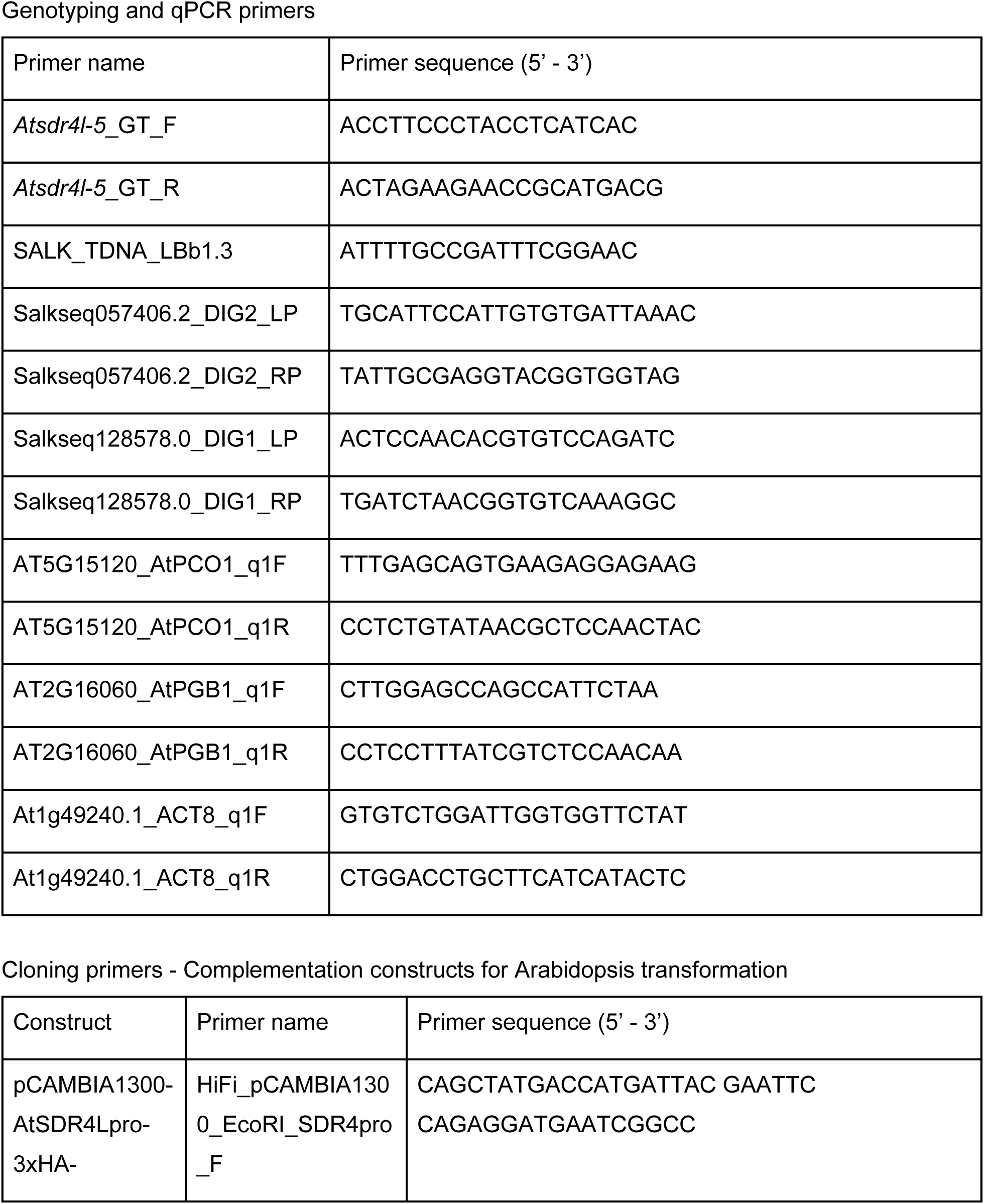
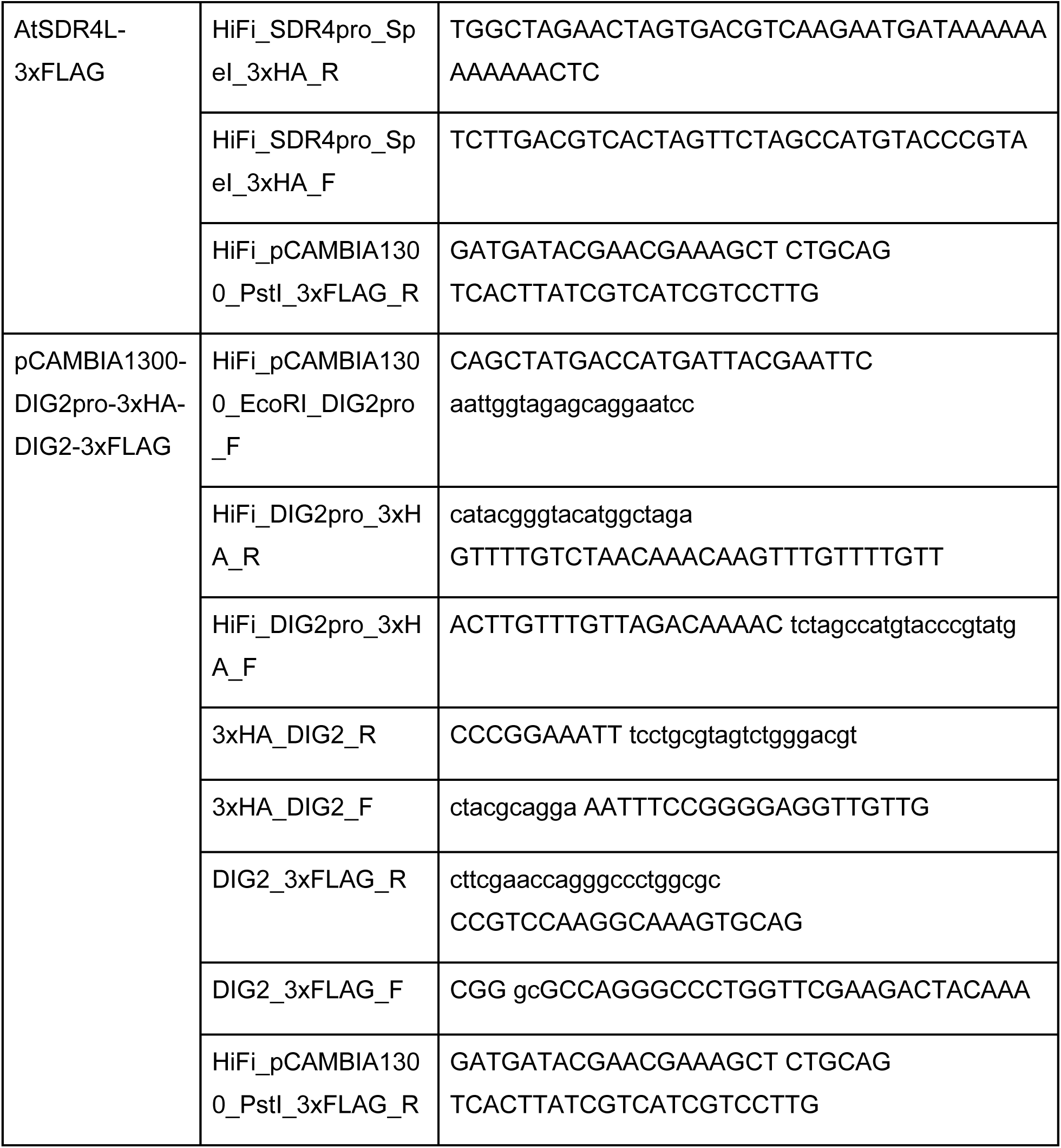
Primers.

**AtSDR4L and paralogs**

AT1G27461 (AtSDR4L/SFL1/ODR1), AT3G48510 (DIG1/SFL2/AITR2), AT5G50360 (DIG2/SFL4/AITR5)

**Selected genes in the main figures**

AT2G27380 (EPR1), AT3G24650 (ABI3), AT1G18100 (MFT), AT3G26790 (FUS3), AT5G45830 (DOG1), AT3G24220 (NCED6), AT3G14440 (NCED3), AT2G29090 (CYP707A2), AT3G03450 (RGL2), AT2G14210 (AGL44), AT2G36270 (ABI5), AT4G13260 (YUC2), AT3G44310 (NIT1), AT3G44300 (NIT2), AT1G08980 (AMI1), AT1G01480 (ACS2), AT4G11280 (ACS6), AT3G23150 (ETR2), AT5G03280 (EIN2), AT1G77120 (ADH1), AT3G03270 (HRU1), AT4G33070 (PDC1), AT5G15120 (PCO1), AT5G39890 (PCO2), AT2G16060 (PGB1), AT1G02400 (GA2OX6), AT3G03450 (RGL2), AT5G17490 (RGL3), AT2G37640 (EXP3) AT2G40610 (EXP8), AT5G17430 (BBM), AT4G37940 (AGL21), AT1G49240 (ACT8)

## Notes

### Competing Interest Statement

The authors have declared no competing interest.

### Summary of Updates

Several main and supplementary figures have been updated in this version.

